# EpImAge: An Epigenetic-Immune Clock for Disease-Associated Biological Aging

**DOI:** 10.1101/2025.03.11.642648

**Authors:** Alena Kalyakulina, Igor Yusipov, Arseniy Trukhanov, Claudio Franceschi, Alexey Moskalev, Mikhail Ivanchenko

## Abstract

**Background:** We present EpImAge, an explainable deep learning tool that integrates epigenetic and immunological markers to create a highly accurate, disease-sensitive biological age predictor. This novel approach bridges two key hallmarks of aging - epigenetic alterations and immunosenescence.

**Methods:** First, epigenetic and immunologic data from the same participants was used for AI models predicting levels of 24 cytokines from blood DNA methylation. Second, open-source epigenetic data (25 thousand samples) was used for generating synthetic immunological biomarkers and training an age estimation model.

**Results:** Using state-of-the-art deep neural networks optimized for tabular data analysis, EpImAge achieves competitive performance metrics against 33 epigenetic clock models, including an overall mean absolute error of 7 years and a Pearson correlation of 0.85 in healthy controls, while demonstrating robust sensitivity across multiple disease categories. Explainable AI revealed the contribution of each immunological feature to the age prediction.

**Conclusions:** The sensitivity to multiple diseases due to combining immunologic and epigenetic profiles is promising for both research and clinical applications. EpImAge is released as an easy-to-use web tool that generates the age estimates and levels of immunological parameters for methylation data, with the detailed report on the contribution of input variables to the model output for each sample.

## 1. Background

Aging is characterized by progressive deterioration of cellular, tissue, physiological and cognitive functions, manifesting through multiple interconnected biological mechanisms that collectively impact organismal health and longevity [1, 2]. It is followed by an increased susceptibility to age-associated pathologies, including cancer, metabolic disorders, cardiovascular diseases, and neurodegenerative conditions [3–6]. According to a broadly accepted classification, there are nine basic hallmarks of aging: genomic instability, telomere attrition, epigenetic alterations, loss of proteostasis, deregulated nutrient sensing, mitochondrial dysfunction, cellular senescence, stem cell exhaustion, and altered intercellular communication [4]. While their interconnectedness is beyond doubt, disentangling it still remains a challenge. Here we aim to advance bridging the age-related epigenetic modifications and immunosenescence.

One of the most frequently addressed epigenetic mechanisms of aging is DNA methylation, which can lead to genomic instability, disrupting gene regulation and cellular processes, and contribute to the development of age-related diseases [1, 7, 8]. Age-related changes are also linked to immune system remodeling [9–11], the development of persistent inflammation, and the associated imbalance between pro- and anti-inflammatory mechanisms [12, 13]. This underlies the concept of inflammaging, a major paradigm of immunosenescence [14]. Age-related epigenetic remodeling is closely linked to inflammaging: epigenetic alterations contribute to chronic inflammation and impaired anti-inflammatory processes [15–18], inflammation promotes heterochromatin loss and age-related hypomethylation [19–21].

Aging clock models are widely used to estimate an individual’s biological age, which reflects the current health status and the risk of developing age-related diseases. Among them, epigenetic clocks receive much attention due to the good standardization and accumulating availability of DNA methylation data, and the model development has surpassed several generations by now [22–28]. Such models have also been studied in the context of sensitivity to different disease groups (cardiovascular, neurodegenerative, oncological, endocrine, psychiatric, viral), namely, by the ability to detect accelerated aging in patients [29–31]. The clocks that characterize aging of the immune system are much less developed. Current models utilize different sets of markers (various plasma proteins, including cytokines and chemokines) to estimate age and are sensitive to certain age-related pathologies, including cardiac, renal, and neurodegenerative diseases [32–37]. The acceleration of epigenetic and inflammatory aging (defined as a discrepancy between biological age and chronological age) is influenced by a multitude of factors, including, but not limited to, diseases and pathological conditions, sex, race/ethnicity, environment, socioeconomic status, stress and others [7, 29, 38–40].

The convergence of two hallmarks of aging, epigenetic modifications and immunosenescence, can be investigated by a simultaneous analysis of the two profiles: epigenetic and immunological. However, the limited prevalence of immunological data represents a significant challenge. A possible solution could make use of the recently proposed methodology of estimating the levels of individual blood plasma proteins from DNA methylation data, with a further objective of mortality prediction [41, 42]. This approach can also underpin the construction of aging clock models and uncovering associations with various pathologies.

In this paper we present an AI-based approach that infers the inflammatory profile (cytokines and chemokines levels) from DNA methylation data, and makes use of it for the epigenetic-immune clock model (EpImAge). Epigenetic data from more than 25000 open-access samples were used for developing the age predictor model. We performed an extensive comparison of EpImAge to 33 existing epigenetic age models both in terms of age prediction quality metrics (mean absolute error, correlation coefficient) for healthy samples and sensitivity to different diseases (covering 13 chapters of ICD-11). The impact of individual immunologic parameters on age prediction was analyzed using explainable AI approaches.

For ease of application, the proposed model is published together with a user-friendly web interface, which only requires uploading a table with methylation data and age. The interface is able to handle the missing values in the data by providing options for imputation, which together with estimation of immunological parameters provides a significant added value to epigenetic age estimation. The web interface also produces plots of predicted age as a function of chronological age and the distribution of age acceleration. Detailed explanation of the effect of each feature on the age estimate and the significance of each immuno-marker in a particular sample compared with the overall distribution of similarly aged individuals is also implemented. The web interface is openly available from the HuggingFace platform [43].

## 2. Methods

### 2.1. Data Collection and Processing

#### 2.1.1. Concurrent data from immunologic and epigenetic profiles

Initial model development utilized a proprietary dataset comprising paired immunological and epigenetic profiles collected between 2019-2023, encompassing both healthy controls and disease states. It included: healthy samples from Nizhny Novgorod region (139 participants aged 20 to 101 years), healthy samples from Yakutia (84 participants aged 19 to 99 years), samples with ESRD from Nizhny Novgorod region enrolled by “FESPHARM NN” hemodialysis centers (106 participants aged 25 to 88 years). The peculiarities of the study procedure, all possible inconveniences and risks were explained to all participants. Each participant filled in the consent for personal data processing and signed an informed consent taking into account the principle of confidentiality (availability of personal data only to the research team, presentation of data in a common array). The study was approved by the local ethical committee of Nizhny Novgorod State University. All procedures were in accordance with the Declaration of Helsinki 1964 and its later amendments.

To obtain immunologic profiles, the analysis was performed on plasma using the K3-EDTA anticoagulant, without hemolysis and lipemia. Plasma was thawed, spun (3000 rpm, 10 min) to remove debris, and 25 µl was collected in duplicate. Plasma with antibody-immobilized beads was incubated with agitation on a plate shaker overnight (16–18 h) at 2–8°C. The Luminex® assay was run according to the manufacturer’s instructions, using a custom human cytokine 46-plex panel (EMD Millipore Corporation, HCYTA-60 K-PX48). Assay plates were measured using a Magpix (Milliplex MAP). Data acquisition and analysis were done using a standard set of programs MAGPIX®. Data quality was examined based on the following criteria: standard curve for each analyte has a 5P R2 value > 0.95. To pass assay technical quality control, the results for two controls in the kit needed to be within the 95% of CI (confidence interval) provided by the vendor for> 40 of the tested analytes. No further tests were done on samples with results out of range low (< OOR). Samples with results out of range high (> OOR) or greater than the standard curve maximum value (SC max) were not tested at higher dilutions.

To obtain epigenetic profiles, Phenol Chloroform DNA extraction was used. DNA was quantified using the DNA Quantitation Kit Qubit dsDNA BR Assay (Thermo Fisher Scientific), and 250 ng was bisulfite-treated using the EpiMark Bisulfite Conversion Kit (NEB) with case and control samples randomly distributed across arrays. The Illumina Infinium MethylationEPIC BeadChip [44] was used according to the manufacturer’s instructions. DNA methylation is expressed as β values, ranging from 0 for unmethylated to 1 representing complete methylation for each probe. DNAm data preprocessing, normalization, and batch effect correction were performed with the standard pipeline in the ChAMP R package [45]. During preprocessing probes with a detection p value above 0.01 in at least 10% of samples were removed. Functional normalization of raw methylation data was performed using minfi R package function [46].

#### 2.1.2. GEO

We used the R packages GEOmetadb [47], GEOquery [48] to perform a preliminary analysis of all available datasets in the GEO repository [49]. These tools give access to metadata related to samples, platforms, and datasets. The Python package GEOparse [50] was used to retrieve features and fields in the datasets, and in some cases to automatically download pre-processed methylation data.

To train the EpImAge epigenetic-immune clock model, we selected data from healthy controls only. To test the models’ sensitivity to diseases, we selected datasets that contain both cases and controls to examine whether there is age acceleration in patients with diseases compared to healthy controls. All samples are required to have age and sex information. This is necessary for model training as well as for calculating estimates of other epigenetic models and age acceleration. Functional normalization of raw methylation data (if available) was performed using minfi R package function [46].

### 2.2. Model Development and Architecture

#### 2.2.1. Feature selection

The epigenetic data have a huge dimensionality (Illumina 450k and Illumina EPIC standards intersection contains more than 400,000 CpG sites), and only a part of them are associated with certain parameters, so we performed feature selection before model training. Feature selection also helps to reduce model training time, improve quality metrics by reducing the number of noisy features, and reduce the risk of overfitting. At the first step of estimating immunologic feature levels from epigenetic data, we used the mRMR (minimum Redundancy - Maximum Relevance) approach for feature selection, which was first proposed in the context of gene expression data [51]. This approach ranks features based on their importance in predicting the target variable (in our case, immunomarker level), with importance necessarily taking into account redundancy and relevance. The relevance of each individual feature is estimated based on the f-statistics with the target variable, redundancy takes into account the correlation of a new feature with the selected ones in the previous iterations (if a new feature is highly correlated with the already selected ones, it is unlikely to provide much new information). The mRMR approach works iteratively, selecting one new feature at a time. At the first step, 100 CpG sites for each immunomarker were selected using mRMR to build models for estimating immunomarker levels from epigenetic data. A total of 2936 CpG sites were used to estimate all 32 immunologic parameters (all CpGs are listed in Supplementary Table S2).

At the second step of age prediction from immunologic estimates, feature selection is also used. Due to their small number, the Pearson correlation coefficient with a threshold of 0.5 is used here - only immunomarkers estimates correlating with their real values are used for age prediction (for example, GrimAge model uses threshold of 0.35). Thus, 24 immunologic markers are used to estimate age, which, in turn, are estimated from 2228 CpG sites.

#### 2.2.2. Models training

At the first step, a separate model is built for each immunologic parameter. It is important to note that the model does not predict the actual levels of immunologic indicators, but rather the log-transformed values. Transforming the target variable (in this case, immunomarker levels) is often used in data with large variance to reduce skewness, bringing the distribution of the data closer to normal, and improves the stability of gradient flow in backpropagation. This transformation can significantly improve the performance of machine learning models. Original immunomarkers values can be obtained using inverse transformation.

Model development employed contemporary deep neural network architectures optimized for tabular data analysis, including Multi-Layer Perceptron (MLP), Deep Abstract Network (DANet) [52], Feature Tokenizer and Transformer (FT-Transformer) [53], and Gated Adaptive Network for Deep Automated Learning of Features (GANDALF) [54]. Unlike sequential or image data, tabular data lacks inherent spatial or temporal relationships between features, necessitating specialized architectural considerations. Epigenetic and immunologic data are tabular as they are a set of numerical values for each sample. MLP is one of the simplest neural network architectures, consisting of several dense layers. FT-Transformer is an adaptation of the Transformer architecture to tabular data, GANDALF uses a new tabular processing unit Gated Feature Learning Unit (GFLU) with gating mechanism and built-in feature selection, DANet is focused on abstract layers whose main idea is to group correlated features and create higher level abstract features from them.

Before training the models, all data was divided into 2 parts −80% for training/validation and 20% for independent testing. During model training cross validation is used - sequential division of data into training and validation subsets in the ratio of 3 to 1, until all data have been in the role of training and validation. Cross-validation is necessary to determine the best combination of model hyperparameters that provide the best metrics. The final model results were computed on a test subset (not involved in model training at all). To evaluate the quality of the models, we consider the following metrics: MAE (mean absolute error), Pearson correlation coefficient.

The second step uses the same neural network architectures with cross-validation and hyperparametric search. Stratification of samples by age was also used here - in each dataset all samples were divided into 5 quintiles (groups) by age, each group must contain at least 5 samples. If this condition cannot be met (the dataset is too small to provide at least 5 samples in each group), it is completely placed in the test data.

#### 2.2.3. XAI

Deep neural network architectures are often black boxes with non-transparent decision-making principles. This complicates the process of tracking model errors and understanding the reasons why models make certain decisions. However, at present, explainable artificial intelligence approaches are being actively developed that can explain why models predict certain values, in particular, when solving a regression problem. One of the most common approaches are SHAP values [55], which use game theory principles. They show how a particular value of each feature changes the final model prediction for each sample. As a result, they help identify the features that contribute most to the final model age prediction.

### 2.3. Statistical Analysis and Validation

#### 2.3.1. Statistical tests

One of the stages of clock model analysis is to check their association with different diseases. The distributions of age acceleration values between the different considered groups (controls and different types of cases) were tested using the Mann-Whitney U-test [56]. This is a nonparametric test for comparing the results of two independent groups and is used to test the probability that two samples come from the same population, with a two-sided null hypothesis that the two groups are not the same. Statistical significance was assessed using multiple testing corrections with the Benjamini-Hochberg procedure to control the false discovery rate (FDR) at α = 0.05. All reported p-values are FDR-adjusted unless otherwise specified [57].

#### 2.3.2. Epigenetic clock models

The implementations of existing epigenetic clock models against which the EpImAge model was compared were taken from the pyaging library, which aggregates a large number of clock models for different types of data, organisms, and tissues [58]. For inclusion in the current work, we selected human epigenetic clock models that estimate both age and age-associated metrics and can be applied to a wide age range (in particular, clocks that work only for newborns or children, as well as for centenarians, were excluded). A total of 34 models of epigenetic clocks were selected for comparative analysis: Hannum [23], Horvath [22], Lin [59], epiTOC1 (mitotic-like clock, estimates time of cancer) [60], ZhangMortality (estimates mortality risk) [61], DNAmPhenoAge [24], SkinAndBlood [62], GrimAge [25], DNAmTL (estimates telomere length) [63], ZhangEN, ZhangBLUP [64], Han [65], DunedinPACE (estimates pace of aging) [27], AltumAge [66], PCHannum, PCHorvath, PCPhenoAge, HRSInCHPhenoAge, PCSkinAndBlood, PCGrimAge, PCDNAmTL (PC modifications of corresponding epigenetic clock models) [67], GrimAge2 [68], DNAmFitAge [69], ENCen40 [70], YingAdaptAge, YingCausAge, YingDamAge [28], StocH, StocP, StocZ (stochastic versions of Horvath, DNAmPhenoAge, ZhangMortality epigenetic models) [71], stemTOC (stochastic mitotic-like clock, estimates time of cancer) [72], RetroelementAgeV1, RetroelementAgeV2 [73], IntrinClock [74].

#### 2.3.3. GSEA

We merged the CpG sites that were selected for models of immunologic parameters estimation and identified their corresponding genes, 1560 genes in total (Supplementary Table S3). For these genes, we performed gene-set enrichment analysis using the Enrichr library [75–77]. This library contains more than 500 thousand terms from 230 libraries. All libraries in Enrichr are organized into 8 groups according to the topics of gene sets and terms: Transcription, Pathways, Ontologies, Diseases/Drugs, Cell Types, Misc, Legacy, Crowd. Groups Misc (miscellaneous) and Legacy (earlier, already out-of-date versions of libraries) were not considered. We also removed from consideration all libraries and terms related to other organisms (not humans). The resulting list of terms was filtered by adjusted p-value with a threshold of 0.001 and included 314 terms (Supplementary Table S4).

### 2.4. Web interface

The web interface for the EpImAge model is based on the Gradio framework [78], which helps to create user-friendly applications to facilitate the use of machine learning models. The application supports imputation of missing values in the uploaded data (kNN, mean, median), inference of trained models estimating age and immunomarker levels, application of XAI (SHAP) for local explainability of the model result for each sample. To make the model available to a wide community of users, it is hosted on HuggingFace, one of the largest platforms for hosting AI models, datasets and interfaces for them (spaces).

## 3. Results

### 3.1. Study Design and Model Development Pipeline

EpImAge was constructed in two steps. First, we established the correspondence between epigenetic markers and immunological parameters and built a set of AI models that predict the levels of plasma cytokines and chemokines based on DNA methylation data. Second, we produced a two-level biological clock model that employs the AI-predicted levels of immunological parameters as an input for the AI-based age estimate (Figure 1A, 1B).

**Figure 1.**
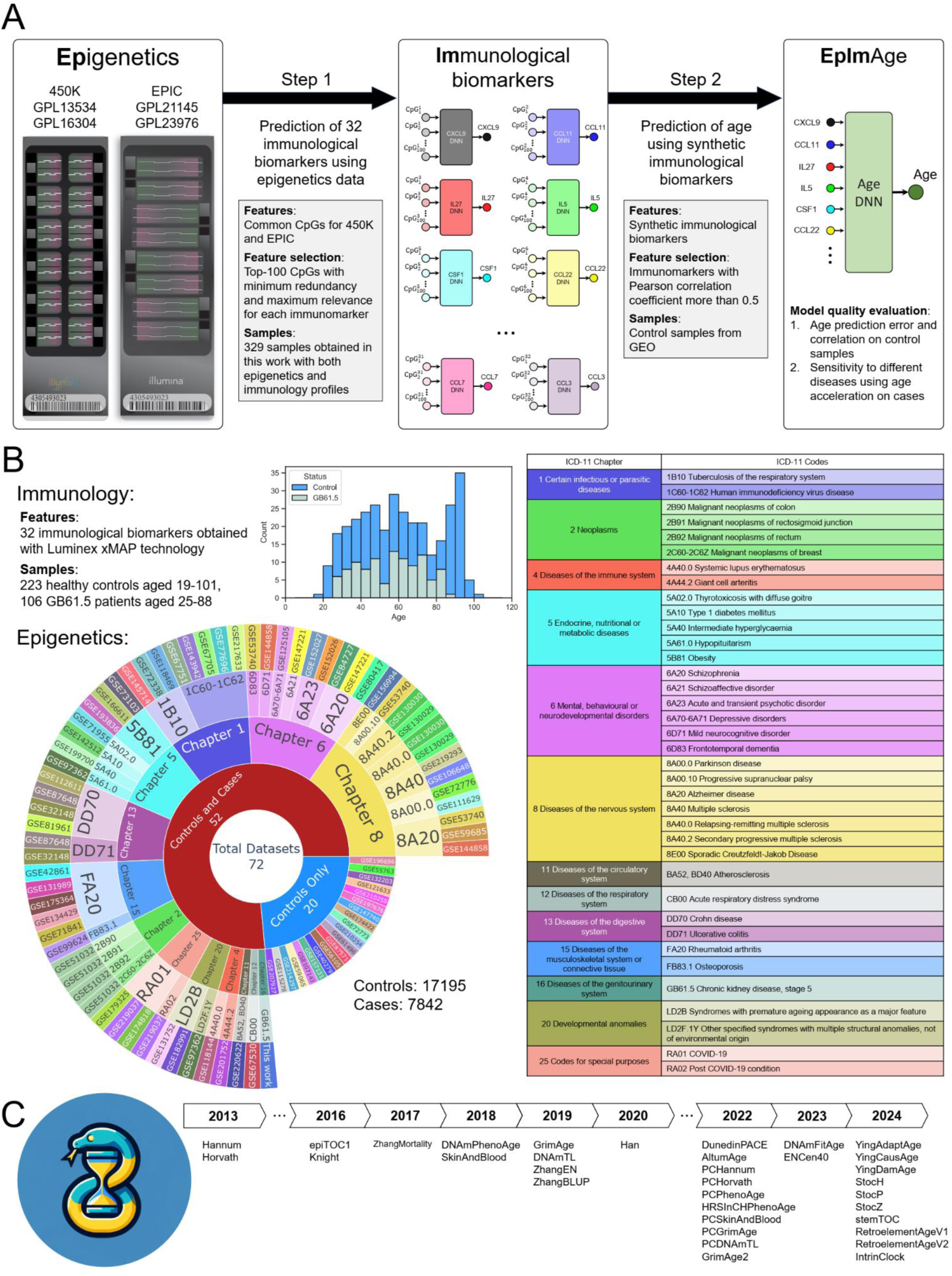
Study design. (A) Main steps in the development of the epigenetic-immune (EpImAge) clock model. First, immunologic markers were estimated from simultaneous epigenetic profile data. In result we obtained 32 neural network models (one for each immunomarker) predicting immunomarker levels. Second, the age estimation model based on synthetic immunologic parameters derived from the open source methylation data, was developed and evaluated against controls and cases groups. (B) Description of data used in this study. The histogram represents the age distribution of the participants whose data were used at the first step. Sunburst diagram characterizes the open source epigenetic data used at the second step: the number of datasets, the number of datasets with controls and cases; ICD-11 chapters and codes for diseases in the studied case groups. The outer level of the diagram gives datasets codes in the NCBI GEO repository. The table on the right accounts for the chapters and particular ICD-11 codes; the samples with these codes were used for the disease-sensitivity test of the resulting clock model. (C) Timeline of development of epigenetic clock models (calculated via pyaging) assessed in our study.

The first step utilized simultaneous epigenetic and immunologic profile data. We make use of the blood sample analysis results for 329 residents of the Russian Federation, aged 19 to 101 years, and recruited in 2019-2023 [33, 36, 79]. Participants included a group of healthy controls (223 participants, 19 to 101 years old) and a group of chronic renal disease patients (106 participants, 25 to 88 years old), the latter broadening the scale of biomarker values, given a certain difference in epigenetic and immunologic profiles between patients and healthy controls [36]. Healthy control group included the representatives from the general population without chronic diseases in the acute phase, cancer, acute respiratory infections, and pregnancy; the chronic renal disease group included patients with stage 5 disease regularly undergoing dialysis. The set of analyzed CpG sites corresponds to the probes common to Illumina 450k and EPIC standards, to ensure a broader applicability of EpImAge, and immunologic profiles are given by levels of 32 cytokines and chemokines.

To estimate immunologic parameters based on epigenetic profiles, first, we used the mRMR (minimum redundancy - maximum relevance) approach for feature selection and reducing redundancy [80]. In result, we obtained the sets of 100 CpG sites for each of the 32 immunomarkers, and found them to have a fairly low overlap (altogether, 2936 different CpG sites were selected to construct predictors for 32 immunomarkers cf. Supplementary Table S1). Next, we constructed 32 separate deep neural networks, one for each immunologic parameter, with 32 different sets of 100 CpG sites serving as inputs. We used the state-of-the art deep neural network architectures for tabular data with training, validation and test subsets division; cross-validation and hyperparametric search were used to select optimal hyperparameter sets. The procedure is illustrated in Figure 1A, Step 1.

The second step addressed age prediction from synthetic immunologic markers (Figure 1A, Step 2). At this stage we made use of public blood DNA methylation datasets from the NCBI GEO repository [49]. We limited our search to Illumina 450k and EPIC standard methylation datasets that have age and sex annotation. Altogether, the data from 17195 healthy participants from 72 datasets with the overall age range starting from a few months to 101 years were used for building the clock. We performed feature selection, retaining only immunologic parameters with the correlation coefficient between real and predicted values greater than 0.5 (see Section 3.2 for details). Synthetic values of immunologic parameter levels we calculated by means of the models developed in the first step. The same model training process was used at this step. Further on, the data from patients with various diseases were used to analyze the sensitivity of the clock model to ICD-11 (see Section 3.4 for details). We also compared the performance of EpImAge with multiple existing epigenetic clocks (Figure 1C).

### 3.2. Predicting immunologic biomarkers from epigenetic data

We start with building deep neural network models that estimate the levels of immunologic markers taking epigenetic data as the input (one neural network - one immunomarker). First, for each immunomarker we selected top 100 CpG sites according to mRMR approach (cf. listed results in Supplementary Table S2) and investigated corresponding correlation coefficients between levels of immunomarkers and their CpGs (represented as a heatmap at Figure 2A). In general, the absolute values of the correlation coefficient did not exceed 0.6 for the best CpG sites, and remained close to 0 for CpG sites at the end of the first hundred. Correlations were not strong for many immunomarkers even for the top CpG sites, the best results were observed for CXCL9, CCL11, IL1B, IL6, GCSF.

**Figure 2.**
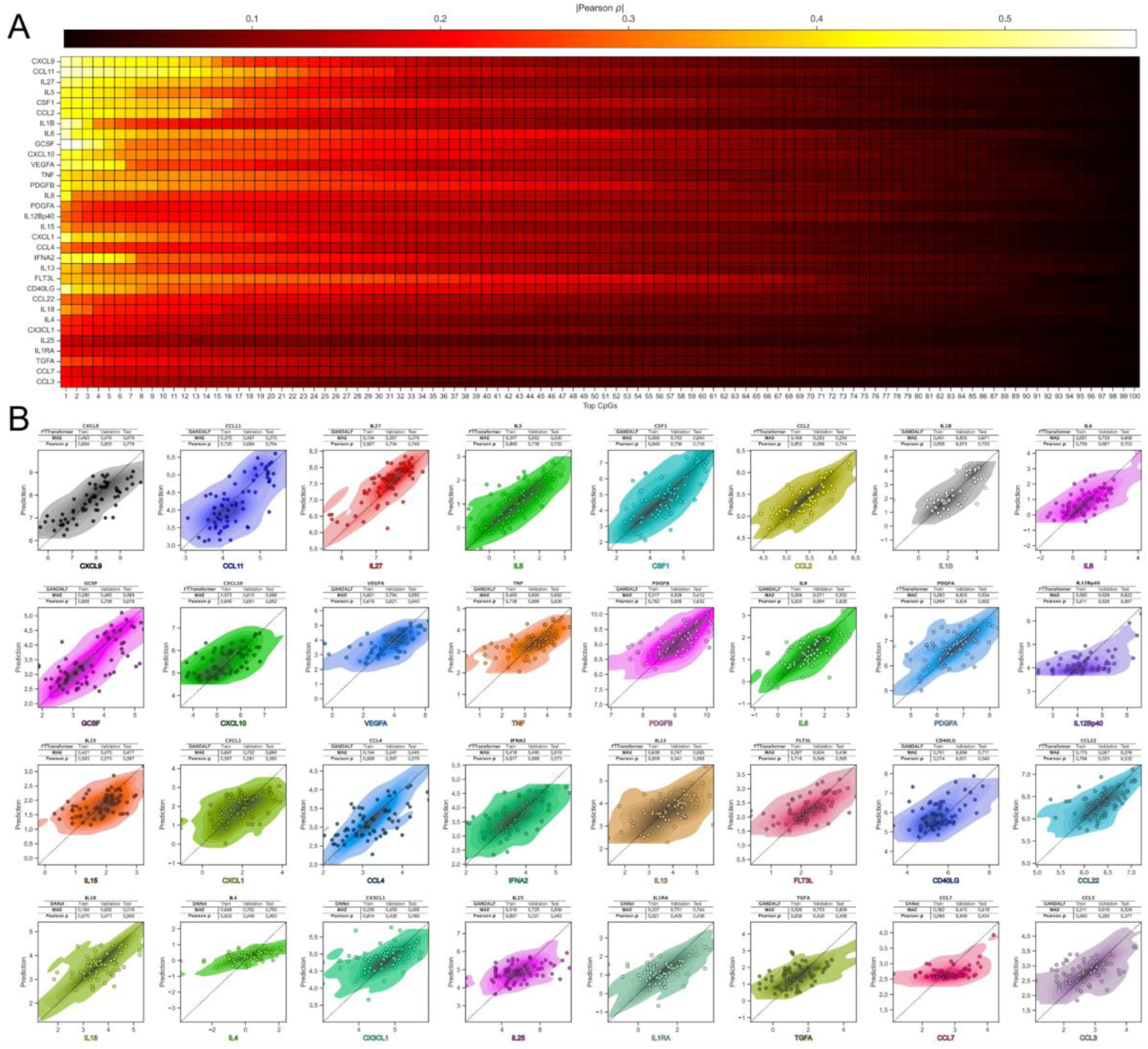
Results of the evaluation of immunologic parameters based on DNA methylation data. (A) Correlation map representing the relationship between immunologic parameters and top 100 CpG sites for each. The absolute value of the Pearson correlation coefficient is shown in color. (B) Results of neural network models for all immunomarkers. For each immunomarker, the relationship between real and predicted levels is presented (logarithmic scale), KDE corresponds to train-validation data, scatter corresponds to test data. The table for each immunomarker shows MAE values and Pearson correlation coefficient for train, validation, test data separately, and the architecture for the best final model is also presented.

Next, we trained individual neural networks for different immunomarkers based on specific top 100 CpG sites. In Figure 2B we illustrate the relationships between the real and predicted values (in logarithmic scale) and annotate the main characteristics of each model (best model architecture, MAE and correlation coefficient for all subsets). The results indicate varying quality of model performance in individual immunomarker predictions. The strongest prediction accuracy is achieved for CXCL9 with a correlation coefficient of 0.78 in test data, while the worst result for CCL3 was r=0.37 only. Nevertheless, 24 of the 32 analyzed immunomarkers manifest correlation coefficients exceeding 0.5 (on test), establishing their reliability for downstream age prediction.

We identified the genes for the CpG sites participating in immunomarker prediction models, (Supplementary Table S3) and performed the Gene-Set Enrichment Analysis (GSEA). We used Enrichr, which aggregates 228 different libraries with lists of genes corresponding to different structures and processes in the human body [75–77]. The resulting list of all significant terms that were found in the libraries is given in Supplementary Table S4. The most common and comprehensive term libraries are Gene Ontology (GO) [81] and KEGG [82], so we considered the significant terms (with adjusted p-value < 0.05) from these libraries separately (Figure 3A). In the GO Biological Process library, the terms related to junction organization and assembly (cell junction assembly and adherence junction organization), cell adhesion (calcium-dependent cell-cell adhesion via plasma membrane cell adhesion molecules and cell-cell adhesion via plasma-membrane cell adhesion molecules), as well as aortic morphogenesis and nervous system development were found to be significant. Axon, excitatory synapse and neuron projection terms found in GO Cellular Component are also relevant. Many terms related to cellular compounds can be associated with aging as well as with the activation and functioning of the immune system during the onset and spread of various diseases, and also during wound healing [83–87]. The functioning of the nervous system declines with age, including degeneration of axons and synapses, which is associated, among other things, with the development of age-associated neurodegenerative diseases [88, 89]. The only significant term in the KEGG library concerned phagocytosis, a process that is actively involved in host-defense mechanisms, through which cells recognize, engulf and digest extrinsic particles (bacteria, apoptotic cells, cellular debris and others) [90]. Interestingly, phagocytosis being closely related to the functioning of the immune system and maintenance of homeostasis, is subject to changes during aging, and phagocyte dysfunction with age may lead to the development of age-associated diseases [91]. Some noteworthy terms from the Enrichr-based list include: epigenetic rejuvenation of mesenchymal stromal cells derived from induced pluripotent stem cells [92]; NF90 target proteins IRF3 and IRF9, crucial regulators of the interferon pathway involved in immune response [93]; pioneer transcription factor ASCL1 regulating neurogenesis and chromatin remodeling [94], and age-associated changes in lncRNA expression and immune system processes [95]. We additionally visualize the results by the word cloud, the word size corresponding to the frequency of the keywords from the papers that investigate the found terms (Figure 3B). Immune activation, cancer (and different individual types of cancer), histone modification, blood pressure, and embryonic stem cells proved to be the most common. It should be pointed out that both aging-related tags and those related to immune function and disease emerged in this word cloud, which suggests a potential interconnectedness in the context of epigenetic and inflammatory aging.

**Figure 3.**
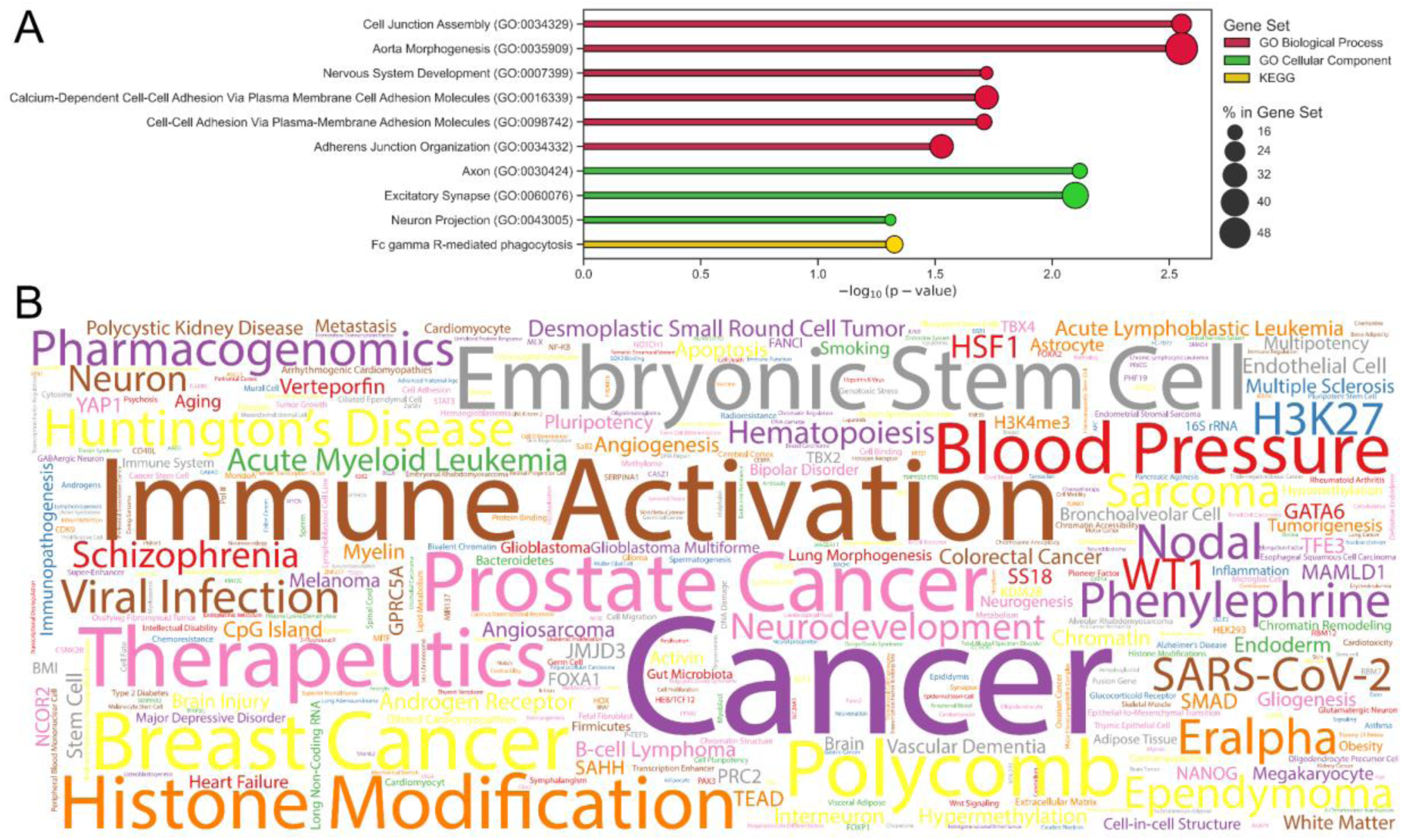
GSEA results for the list of genes involved in the development of immunologic parameter estimation models. (A) GSEA results for GO Biological Process (red), GO Cellular Component (green), KEGG (yellow) libraries. The plot shows the adjusted p-value on a logarithmic scale. The size of the circle for each term shows the percentage of coverage of the gene list corresponding to the term. (B) Word cloud demonstrating the keywords of the papers corresponding to the most significant terms. The size of words corresponds to the frequency of occurrence: the larger the word, the more papers corresponding to the terms have it as a keyword.

### 3.3. EpImAge: epigenetic-immune clock model

Figure 4 summarizes the main characteristics of the EpImAge model, which predicts age from epigenetic data with an intermediate embedding in the form of synthetic immunologic parameters, thereby reflecting both epigenetic and immunologic profiles. The resulting model is based on the DANet deep neural network architecture and demonstrates a good performance with the MAE of 7 years, and correlation between the real and predicted age of 0.85 (the detailed results for train, validation and test are in Figure 4A, all samples participated in model construction are listed in Supplementary Table S5). It is worth noting that the model demonstrates a low bias, partly due to the fact that a modern model, specialized for tabular data, is used (linear models, widely used for epigenetic clocks, yield considerable systematic errors in predicted age [39]). Epigenetic profiles of more than 17k samples with a very wide age range (0-101 years) were used to build the model. The use of such a large number of training data of different standards aimed to increase the robustness of the results, smooth out the batch effects of methylation data from the conditions of their collection and processing, and familiarize the model with the wide variability of epigenetic data. Violin plots for age acceleration values for train, validation, and test data are similar and broad enough to suggest that the model is not overfitted on train.

**Figure 4.**
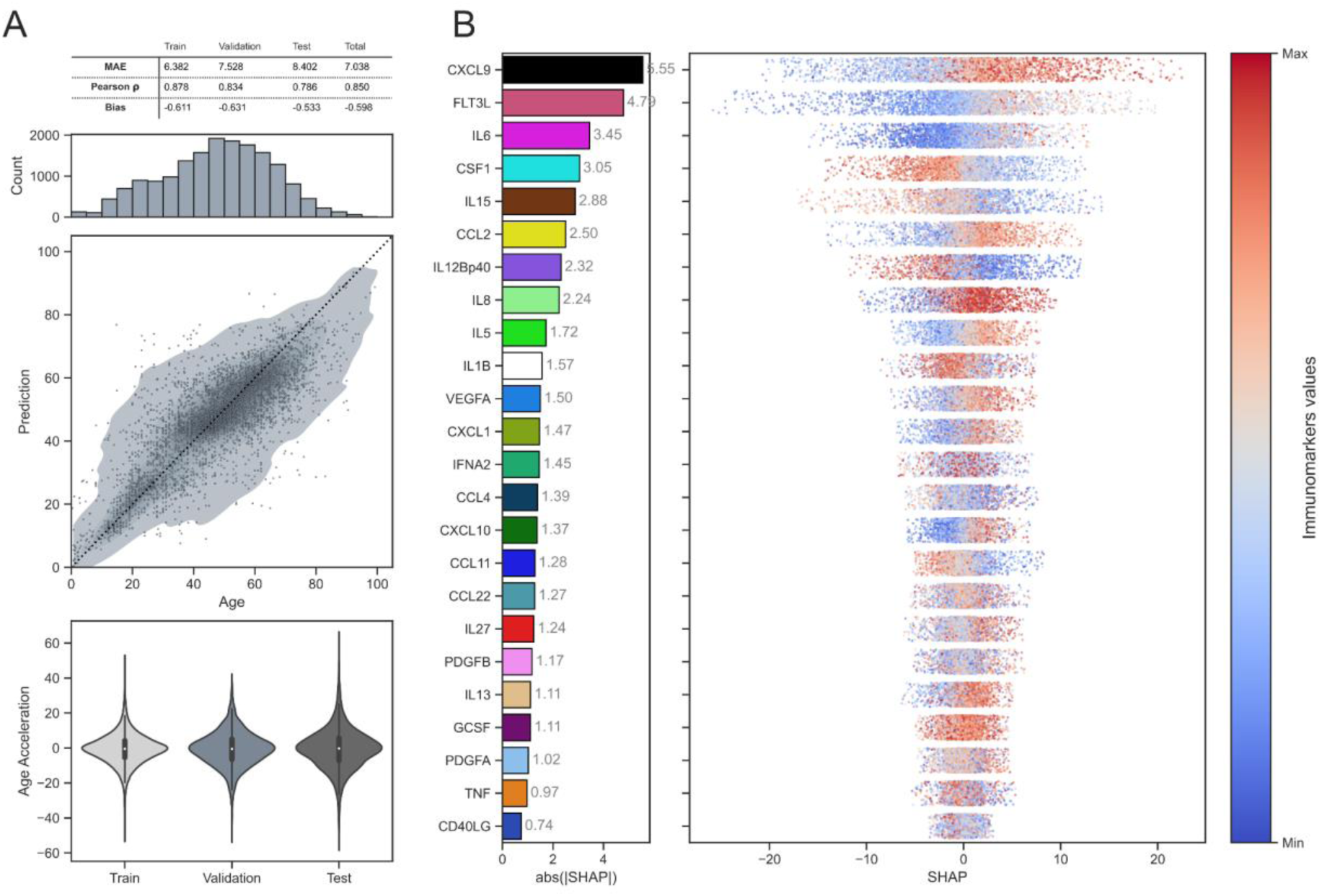
Results of EpImAge epigenetic-immune clock model. (A) (1st row) Main metrics (MAE, Pearson correlation coefficient, bias) total, and separately for training, validation, and test subsets. (2nd row) Histogram of the age distribution of samples used for model training, validation, and testing. (3rd row) Dependence of the predicted age on the real age for the training and validation subsets (gray KDE), for the test subset (gray scatter). The black dashed line corresponds to the bisector. (4th row) Violin plots of age acceleration value distributions for train, validation, and test data. (B) Applying XAI to the EpImAge model. (left) Bar plot illustrating the absolute SHAP values for all immunologic parameters in descending order. (right) Illustration of associations between immunomarkers values and SHAP values. Higher values of each immunomarker are in red, lower values in blue.

Figure 4B shows the results of applying explainable AI approaches to the EpImAge model. The features (synthetic immunologic parameters) are ranked according to their influence on the model’s final decisions as a whole, measured by SHAP values (so-called global explainability). The largest contribution comes from CXCL9, which agrees with previous results [32, 33]. Higher values of CXCL9 correspond to a positive contribution to age estimation, while lower values correspond to a negative contribution. Similar associations of immunologic parameters with age are observed for FLT3L and IL6, while for CSF1 and IL15 decreasing levels are associated with increasing age. The associations of immunologic parameters with age and diseases identified by XAI are described in more detail in Section 3.6.

The performance of the model for individual datasets is detailed presented in Supplementary Figure S1, for the majority of datasets characterized by a wide age range, the total MAE was in the range of 6-8 years.

EpImAge is quite competitive to existing immunologic clock models, although it relies on predicted immunological parameters, rather than directly measured. The iAge inflammatory clock has a higher average error of 15.2 years [32], the chronic renal disease-sensitive ipAGE clock model has a comparable mean absolute error of 7.27 years [36], the small immunologic clock model SImAge also has a comparable mean absolute error of 6.94 years [33], the CyClo age-related brain atrophy-sensitive clock model showed a slightly lower mean absolute error of 6 years, but has also a lower correlation between the real and predicted age of about 0.5 [37]. EpImAge shows comparable results for immunomarkers estimates, working originally on epigenetic data, thus bringing epigenetic and immune components of aging together.

So far, we evaluated the quality of the model’s performance only on healthy samples, without chronic or terminal diseases. However, it is challenging to assess its ability to detect age acceleration in patients with age-associated diseases and other pathological conditions.

### 3.4. Sensitivity of EpImAge and other epigenetic clocks to different diseases

An important property of aging clock models (epigenetic, immunologic and of another nature) is the sensitivity to diseases, the ability to detect age acceleration in patients with various pathologic conditions in comparison to healthy controls.

The relevant datasets from NCBI GEO are shown in Figure 1B. At this stage we selected 52 out of 72, which contain both cases and controls, for the comparison to be possible. We do not consider cases-only datasets due to the heterogeneity of diseases and the inability to compare age acceleration with healthy controls under the same data collection conditions. The datasets give a fairly broad representation by disease group, 13 ICD-11 chapters were found, the number of individual disease codes within each chapter varied from 1 to 7. The outer ring of the chart shows which datasets contain samples with particular diseases (Figure 1B).

Figure 5 summarizes the results of the age acceleration analysis of the EpImAge model for controls and cases, grouped by ICD-11 chapters. The comparison of the mean levels of age acceleration was made using the pairwise Mann-Whitney U-test with a significance threshold of 0.05. Since this test does not recognize the signature of the age acceleration, we additionally take into account bias values when comparing controls and cases. For the infectious and parasitic diseases (Chapter 1), the model detects HIV and HIV-related groups quite well, showing an increase in age acceleration in patients compared to healthy controls (however, there are a few small datasets in which statistical significance is not observed, possibly due to small sample size or dataset specificity). Diseases of the immune system (Chapter 4) are poorly represented in the data; however, there was significant age acceleration in patients with systemic lupus erythematosus. Interestingly, mental, behavioral, and neurodevelopmental disorders (Chapter 6) were detected quite well by the model: for almost all datasets, statistically significant age acceleration was observed in patients with schizophrenia, psychosis, and depression. Noteworthy, the resulting age acceleration can yield different results for the same disease in different datasets (cf. HIV, Crohn’s disease). It could be the consequence of heterogeneity of datasets in question, possibly due to their own batch effects and conditions of data collection and processing.

**Figure 5.**
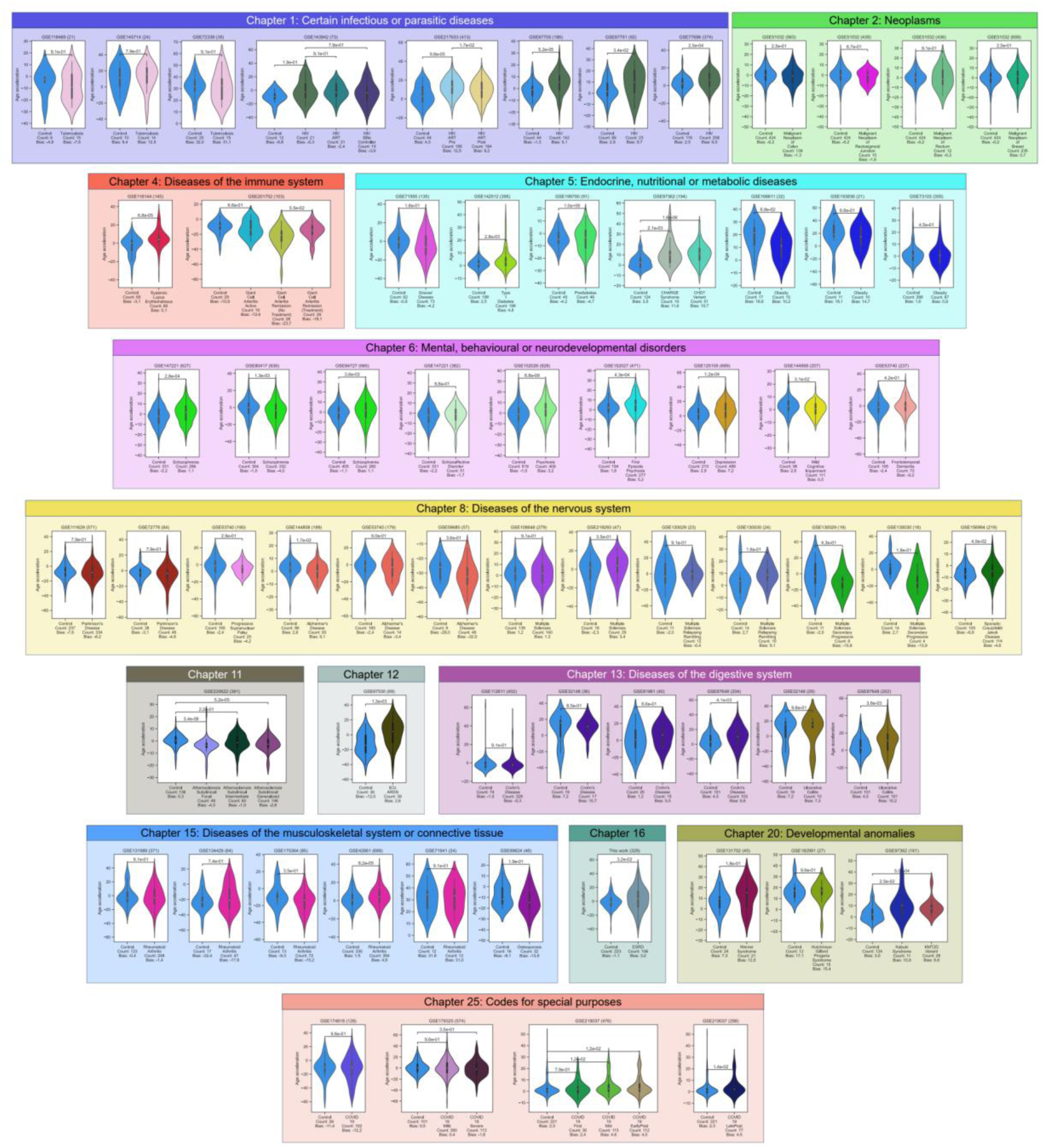
Age acceleration estimated by the EpImAge model for controls and cases for different disease groups. Statistical significance of the difference in mean values of age acceleration was assessed using the pairwise Mann-Whitney U-test with an FDR-adjusted p-value threshold of 0.05 (76 tests in total). Different backgrounds correspond to different chapters of ICD-11. Each plot shows the violins of age acceleration in different groups, with the name of the corresponding dataset (GEO code) and the number of samples in it. The name of the group (control - healthy participants, or cases - patients with different diseases), the number of samples in the group and bias value are given under each violin. P-values are given between the violins.

Next, we performed a large-scale comparison of EpImAge against 33 epigenetic clock models, shown with the development timeline in Figure 1C (for details cf. Methods, Section 2.3, model implementations were taken from the pyaging library). Most of the considered epigenetic models estimate age, some of them estimate age-associated metrics (epiTOC1, ZhangMortality, DNAmTL, DunedinPACE, PCDNAmTL, stemTOC). As the benchmarks for age prediction we made use of MAE and correlation coefficient on controls (applicable only to those clocks that estimate age). Disease sensitivity is characterized by an index defined as the number of tests passed out of the total 76 (applicable to all models).

Figure 6 summarizes the results. The best model in terms of quality metrics for healthy controls has proved to be ZhangBLUP (MAE even less than 3 years, with a correlation coefficient over 0.97), however, it displays almost no disease sensitivity (only 9 tests out of 76 passed). The best model in terms of sensitivity to diseases is DunedinPACE (36 tests out of 76), which estimates the rate of aging rather than age. The HRSinCHPhenoAge and PCPhenoAge models (modifications of the DNAmPhenoAge model) perform quite well, they show high levels of age prediction metrics and rather good sensitivity to diseases. Our EpImAge clock shows competitive results with the quality metrics of most popular models, and rather high sensitivity to diseases. Concurrently, it is also able to estimate immunologic markers levels, and interpret age acceleration in the inflammatory perspective. There are disease groups where EpImAge outperforms most of the other clocks, e.g. for metabolic diseases (Chapter 5) only DunedinPACE passes more disease sensitivity tests but does not estimate age; for developmental anomalies (Chapter 20) models with similar disease group performance have worse control group quality metrics.

**Figure 6.**
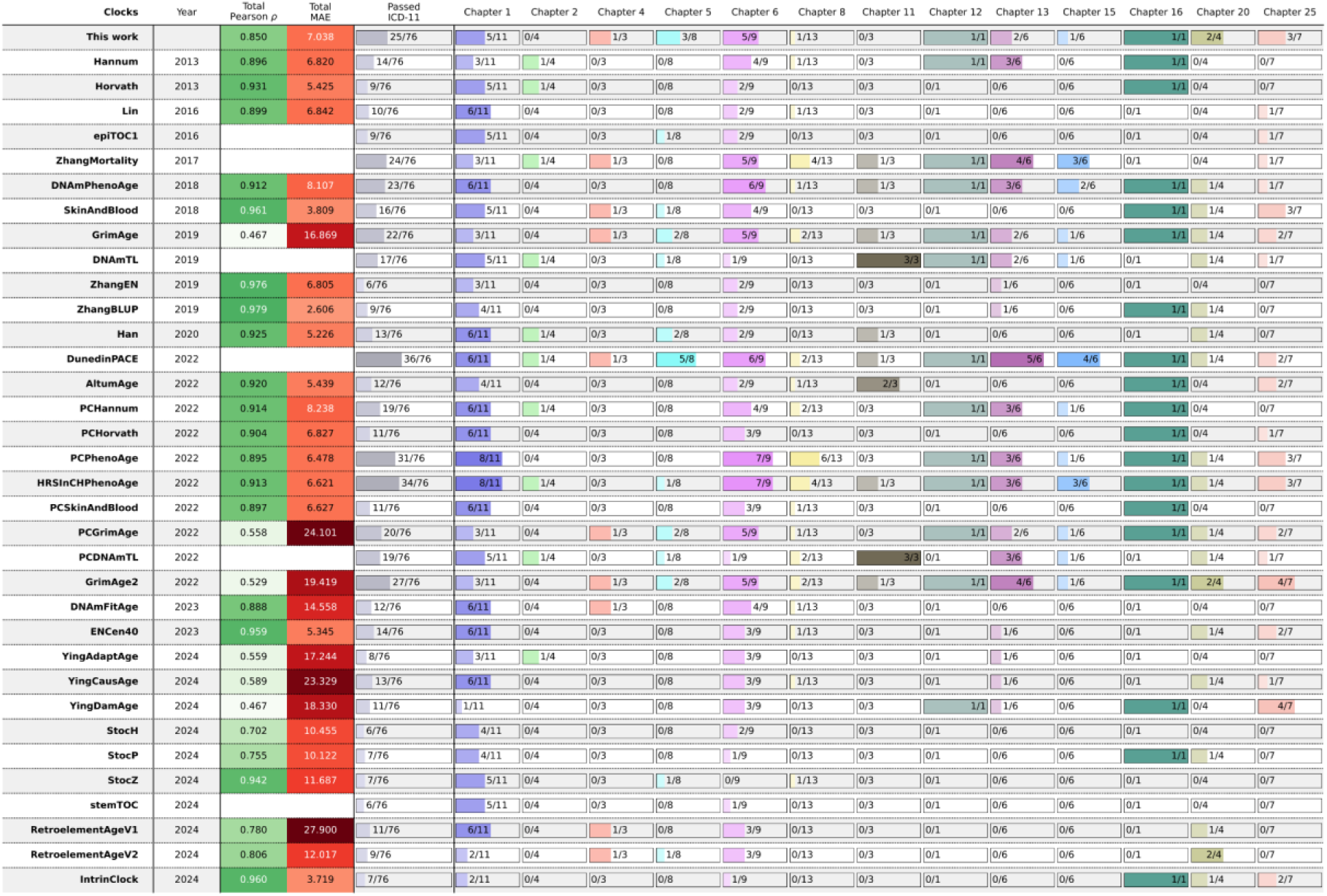
Comparative performance results of different epigenetic clocks (including the ones presented in this paper). The first column gives the names of the clocks. For each epigenetic clock model, the year of development (clocks are ordered by the chronology of their release), common metrics - Pearson correlation coefficient and MAE, and the number of disease sensitivity tests passed (Mann-Whitney p-value < 0.05) are given both overall and for each ICD-11 chapter.

Overall, infectious, mental, respiratory, genitourinary diseases (ICD chapters 1, 6, 12, 16, 25) are detected quite well by a large number of clock models, while neoplasms, immune, nervous, musculoskeletal, developmental diseases (chapters 2, 4, 8, 15, 20) are detected poorly. There are several possible reasons for that. We already pointed out that methylation data can have a strong dependence on the laboratory conditions of the experiment, data preprocessing, and sampling characteristics. Besides, different chapters have different coverage by datasets, infectious, mental, and nervous diseases (chapters 1, 6, and 8) contain 11, 9, and 13 tests, respectively, while respiratory and genitourinary diseases (chapters 12 and 16) contain only 1. Further on, some diseases may not be reflected well by blood DNA methylation. For example, diseases of the nervous system are poorly detected (according to 11 tests). More detailed information on the sensitivity of different models to individual diseases is presented in Supplementary Figure S2, detailed results for all samples are listed in Supplementary Table S5. Thus, different models are good at detecting different groups of diseases and such analysis is useful for selection of a suitable model.

### 3.5. Applying XAI to the EpImAge model

XAI has recently become an integral part of any AI-related research and development. It is a useful and powerful tool in the study of deep neural network models, allowing to open “black boxes”, which in fact are many modern AI models, since the principles of their decision-making are opaque for both developers and users. XAI approaches allow one to explain model behavior from both the global (the general contributions of the model’s features to decision making) and the local (the influence of features on the model’s decision for each particular sample) perspectives, to track and correct model errors. There are many approaches that can be applied to different types of data, specialized for a particular class of models or applied to a wide range of architectures. In this paper we used SHAP, a model-agnostic method based on the game theory ideas that evaluates the contribution of each feature to the change in model prediction (cf. Methods, Section 2.2). For the developed EpImAge model XAI was used to analyze the influence of immunomarkers on relative age acceleration in the analysis of different pathologies, and to specify the particular contribution of each immunomarker, measured in years, to age acceleration.

Figure 7A shows a clustermap illustrating the difference in age acceleration determined by SHAP between cases and controls (positive values correspond to higher values of age acceleration in cases compared to controls, negative values correspond to lower values). The biomarkers in the upper part of the clustermap - CXCL9, IL6, CSF1, CCL4, IL15 - have the greatest total effect on age acceleration. At the same time, some of them consistently contribute to age acceleration for most of the considered diseases (CXCL9, IL6), while some of them contribute to deceleration (CSF1, IL15). Note that there is not a single chapter for which all immunomarkers have only positive or only negative contributions, nor there is a single immunomarker that shows only positive or only negative contributions for all disease chapters. For example, for diseases of the nervous system (Chapter 8), many markers display negative values, while for diseases of the musculoskeletal system (Chapter 15) many markers are positive.

**Figure 7.**
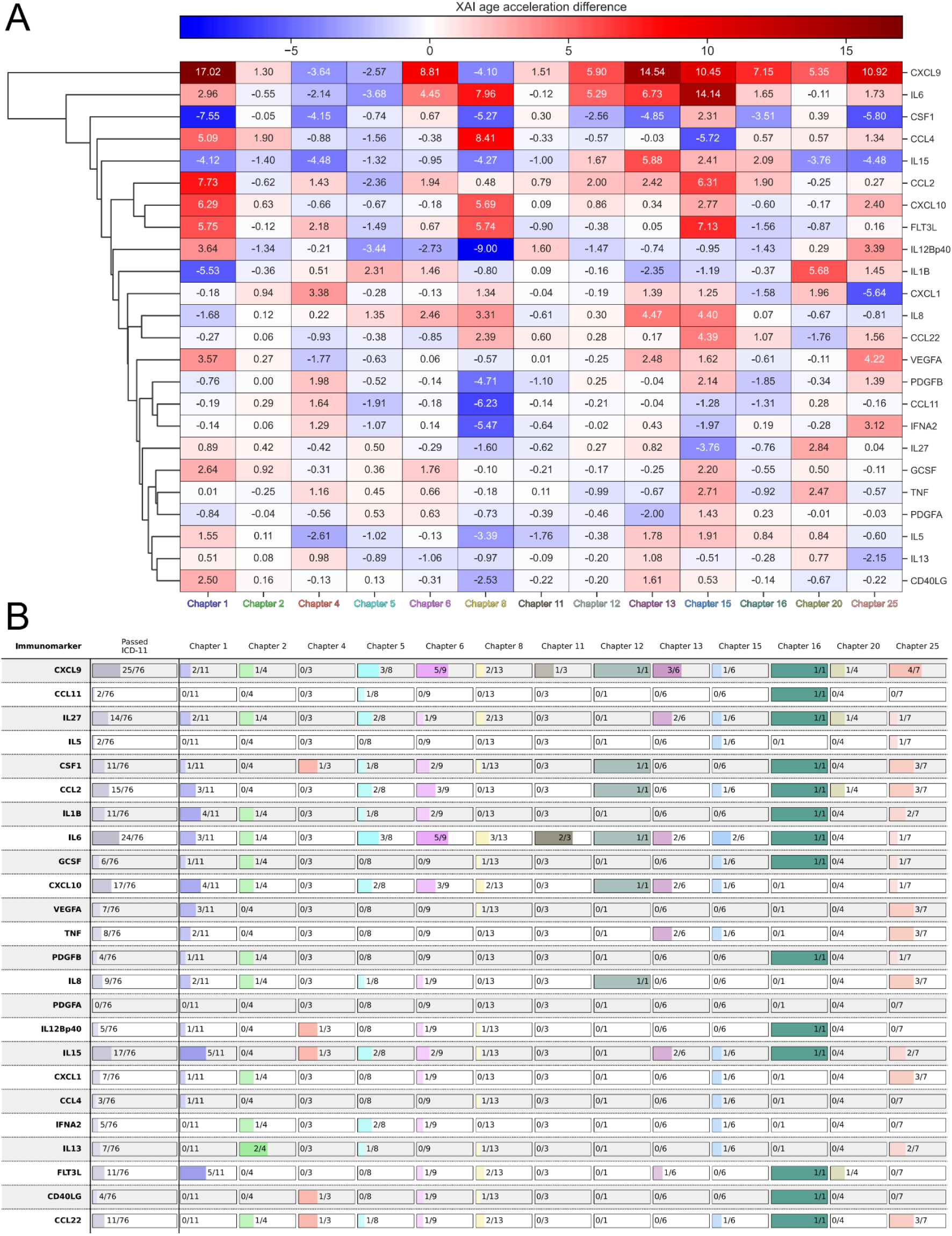
Association of selected immunomarkers with different disease groups. (A) Clustermap illustrating the difference in age acceleration between cases and controls, determined using SHAP values, for all ICD-11 chapters considered. Positive values (red) correspond to higher values of age acceleration in cases compared to controls, negative values (blue) correspond to lower values of age acceleration in cases compared to controls. (B) Comparative results of the association with different disease groups of the immunologic markers studied. For each immunomarker, the number of disease sensitivity tests passed (Mann-Whitney p-value < 0.05) is given both overall and for each ICD-11 chapter.

Together with the results on the difference in age acceleration calculated via XAI, it is interesting to see how individual immunomarkers are associated with different disease groups (Figure 7B). These results agree well with the clustermap, that is CXCL9, IL6, IL15 are most sensitive to diseases. Chapters are detected differently by different immunomarkers, the picture is generally worse than for the clocks. PDGFA does not sense any of the diseases. More detailed information on the sensitivity of different immunomarkers to individual diseases is presented in Supplementary Figure S3.

### 3.6. EpImAge web interface

EpImAge model for epigenetic-immune age estimation is publicly available with a user-friendly interface [43]. The interface takes as input DNA methylation data for 2228 CpG sites (the list can be obtained from the web interface page), and all samples must be assigned with age and have a unique identifier. Functionality allows to process missing values in the data; one of three approaches can be chosen: kNN (k nearest neighbors), mean (filling missing values with mean values for a given CpG), median (filling missing values with median values for a given CpG). The output of the interface provides the overall metrics, MAE and Pearson correlation coefficient, as well as a table with the results of the models: immunomarker estimates, EpImAge and age acceleration values. In addition, interactive scatter and violin plots are provided, illustrating the relationship between chronological age and predicted age, and the distribution of age acceleration values, respectively.

Explainable artificial intelligence for the EpImAge model is also available in the provided web interface. For each individual sample (selected by unique identifier), an interactive waterfall plot can be obtained that shows the contribution of each individual feature to the final model prediction, and how they shift the predicted age relative to the chronological age. For each individual immunomarker, the levels relative to samples of similar age are given (what percentage of samples with similar chronological age have lower levels, and what percentage have higher levels). For top 3 most important immunomarkers, the correspondence age-associated diseases are outlined and supplied with references to literature.

## 4. Discussion

One of the primary objectives of this research was to integrate the two hallmarks of aging, namely epigenetic modifications and immunosenescence. To this end, we conducted a simultaneous examination of DNA methylation data and levels of cytokines and chemokines. We developed models for estimating immunomarker levels from epigenetic profiles and subsequently evaluated their performance on a large cohort of healthy and diseased samples. Estimates of immunologic markers have proven to be most sensitive to infectious and parasitic diseases, including COVID-19, as well as respiratory and genitourinary diseases. The results of studies [96–99] indicate that IL15, FLT3L, CXCL10, IL1B, which demonstrated the most promising outcomes among the estimates, are actually associated with numerous viral, bacterial, parasitic, and fungal infections. Additionally, one of the most extensively studied diseases in recent years, COVID-19 and its associated complications, have been shown to induce alterations in the levels of the majority of the studied immunomarkers [100–107].

CXCL9, CCL2, IL6, CXCL10, and IL8 have been linked to various respiratory diseases [108–112] and retain sensitivity for their level estimates as well. Chronic kidney disease, which has a significant impact on the body’s immune system and is associated with irreversible changes in cytokine levels [36, 113–118], also causes notable alterations in the levels of the majority of synthetic immunomarkers.

The estimates of immunomarkers have proved least sensitive to diseases of the immune, cardiovascular, musculoskeletal systems, and developmental anomalies. This may be attributed to the limited representation of these diseases in the analyzed data. The estimation of immunological parameters is based on a limited number of CpG sites, which may result in the omission of certain aspects of cytokine functionality, which also could be a contributing factor to the reduced sensitivity. Low sensitivity observed with respect to diseases of the nervous system may be attributed, at least in part, to the anatomical location of these diseases, which are frequently confined to the central nervous system and separated from the blood by the blood-brain barrier. Consequently, these diseases may be manifested less conspicuously in blood parameters.

The estimated levels of CXCL9 and IL6 demonstrate sensitivity to almost all disease groups, which is consistent with their important role in the inflammatory response in humans and their association with multiple age-related and pathological changes [100, 108, 113, 119–128]. In conclusion, there is compelling evidence that estimates of immunologic markers mirror associations between their real-world counterparts and various disease groups. However, further investigation is necessary, given the limited scope of diseases represented in the current study.

The use of a large number of epigenetic data for controls and cases allowed us not only to build a model of age estimation and test its sensitivity to diseases, but also to conduct a direct large-scale comparison with other epigenetic clock models. We inferred the consistency of the models and investigated their applicability to different groups of diseases, comparing their performance on cases and controls, confirmed the associations found earlier and identified the new ones.

Clock models that are frequently used in the field have previously been tested on a wide range of data in multiple studies, and in this analysis confirmed associations with a wide range of diseases, to name Hannum [23, 29–31, 129, 130], Horvath [22, 29–31, 36, 129, 131, 132], DNAmPhenoAge [24, 29, 36, 129, 131–134], GrimAge clocks [25, 29, 31, 36, 129, 133–138], as well as DunedinPACE [27, 28, 130, 139–145].

The sensitivity of many clock models to diseases of the immune, digestive, genitourinary systems, and developmental anomalies has been understudied, and this study aims to bridge this gap. The EpImAge model, which combines immune and epigenetic components, and several epigenetic clock models (ZhangMortality and DunedinPACE for arteritis; SkinBlood, both GrimAge, both RetroElementAge for lupus) are able to detect immune system diseases. In the majority of cases, Crohn’s disease and ulcerative colitis (diseases of the digestive system) are either recognized by the models simultaneously or not at all. Instructively, it was found that the chronic kidney disease, that is associated with changes in many immunologic marker estimates, results in significant alterations according to the majority of epigenetic clock models. Furthermore, it appears that profound changes in stage 5 disease are also reflected in the epigenetic profile. The low sensitivity of the clock to developmental abnormalities can be attributed to the rarity of these pathologies, their varied manifestations, and the fact that they may not always align with the genomic regions utilized in the model’s construction.

It is worth noting that the EpImAge model has proved to be the only one to show sensitivity to diabetes, and stands among the best in the sensitivity to a number of other conditions, including systemic lupus erythematosus, psychosis, depression, hypopituitarism, Creutzfeldt-Jakob disease, acute respiratory distress syndrome, chronic renal failure, CHARGE and Kabuki syndromes, COVID-19.

The proposed analysis can be particularly useful in the case of epigenetic clock models that have only recently been developed. There is still a lack of studies on different cohorts, and testing sensitivity to a wide range of diseases is a priority. Noteworthy, the AdaptAge, CausAge, and DamAge models, which are based on a similar principle, exhibit significant differences in their sensitivity to infectious diseases. Stochastic versions of the Horvath, PhenoAge, and Zhang clocks demonstrate varying degrees of sensitivity to mental disorders. Additionally, the RetroElementAge model shows promise in detecting atypical aging patterns associated with developmental abnormalities.

Despite EpImAge’s strong performance, several key limitations must be considered when interpreting results and planning future applications:

Sample Size Constraints: The immunological embedding is based on a relatively modest sample size (n=329), which may limit the model’s ability to capture the full spectrum of immune profile variations.

Data Heterogeneity: Epigenetic data from different datasets exhibit substantial technical variation and batch effects, potentially impacting model generalizability.

Disease Representation: The availability of certain disease groups in public repositories is limited, affecting our ability to comprehensively assess model performance across all pathological conditions.

Feature Selection Methodology: While our mRMR-based approach proved effective, alternative feature selection strategies might yield different or potentially improved marker sets.

Model Architecture Choices: Though we employed state-of-the-art deep learning architectures, the rapidly evolving nature of AI suggests that newer approaches may offer further improvements.

Future work should address these limitations through larger-scale validation studies and exploration of emerging methodologies.

Furthermore, while our validation strategy incorporated multiple independent cohorts, future studies should assess EpImAge’s performance in more diverse populations and clinical settings. Additionally, the relationship between synthetic immunological markers and their biological counterparts warrants further investigation through longitudinal studies. To ensure robust validation, we implemented a multi-tiered strategy: (1) internal cross-validation during model development, (2) held-out test set validation for performance assessment, and (3) external validation using independent cohorts to evaluate the potential to generalization. This comprehensive approach provides confidence in the model’s real-world applicability while identifying potential limitations. EpImAge represents a significant advance in biological age prediction by bridging epigenetic and immunological markers through deep learning. The model’s ability to generate interpretable results while maintaining high accuracy across diverse populations and disease states suggests potential applications in both research and clinical settings. Future work should focus on expanding the validation cohorts, investigating causal relationships between synthetic and biological markers, and developing standardized protocols for clinical implementation.

## 5. Conclusions

We introduced EpImAge, a novel deep learning framework that bridges epigenetic and immunological aspects of aging. Our results demonstrate three key advances: (1) successful prediction of immunological markers from DNA methylation data, (2) accurate age estimation using synthetic immunological profiles, and (3) robust disease sensitivity across multiple pathological conditions. These findings have important implications for both aging research and clinical applications. The model is accessible via a web interface, enabling users to retrieve results (immunologic parameter levels and age estimates) with minimal delay and perform XAI interpretation for each sample.

## List of abbreviations

AI: Artificial Intelligence
COPD: Chronic Obstructive Pulmonary Disease
COVID-19: Coronavirus Disease 2019
DANet: Deep Abstract Network
DNA: Deoxyribonucleic Acid
FT-Transformer: Feature Tokenizer and Transformer
GANDALF: Gated Adaptive Network for Deep Automated Learning of Features
GEO: Gene Expression Omnibus
GFLU: Gated Feature Learning Unit
GO: Gene Ontology
GSEA: Gene-Set Enrichment Analysis
ICD: International Classification of Diseases
HIV: Human Immunodeficiency Virus
KEGG: Kyoto Encyclopedia of Genes and Genomes
kNN: k Nearest Neighbors
MAE: Mean Absolute Error
mRMR: Minimum Redundancy, Maximum Relevance
RNA: Ribonucleic Acid
SHAP: SHapley Additive exPlanations
XAI: eXplainable Artificial Intelligence.

## Data availability statement

The data used to construct the models are included in this published article and its Supplementary Tables. Open-access DNA methylation data were downloaded from the NCBI GEO repository, all corresponding GSE codes are summarized in Figure 1 and GSM codes are in Supplementary Table S5.

## Code availability statement

The source code for the pipeline presented in this paper is publicly available in the repository: https://github.com/GillianGrayson/EpImAge. The web interface for using the EpImAge model is available at: https://huggingface.co/spaces/UNNAILab/EpImAge.

## Authors’ contributions

Conceptualization: AK, IY, MI; Formal analysis: AK, IY; Methodology: AK, IY, MI; Software: AK, IY; Supervision: AT, CF, AM, MI; Visualization: AK, IY; Writing – original draft: AK, IY; Writing – review and editing: AT, AK, IY, CF, AM, MI. All authors contributed to the article and approved the submitted version.

## References

1. Pereira B, Correia FP, Alves IA, Costa M, Gameiro M, Martins AP, Saraiva JA (2024) Epigenetic reprogramming as a key to reverse ageing and increase longevity. Ageing Research Reviews 95:102204. 10.1016/j.arr.2024.102204

2. Guo J, Huang X, Dou L, Yan M, Shen T, Tang W, Li J (2022) Aging and aging-related diseases: from molecular mechanisms to interventions and treatments. Sig Transduct Target Ther 7:1–40. 10.1038/s41392-022-01251-0

3. Wang K, Liu H, Hu Q, Wang L, Liu J, Zheng Z, Zhang W, Ren J, Zhu F, Liu G-H (2022) Epigenetic regulation of aging: implications for interventions of aging and diseases. Sig Transduct Target Ther 7:1–22. 10.1038/s41392-022-01211-8

4. López-Otín C, Blasco MA, Partridge L, Serrano M, Kroemer G (2013) The Hallmarks of Aging. Cell 153:1194–1217. 10.1016/j.cell.2013.05.039

5. Pal S, Tyler JK (2016) Epigenetics and aging. Sci Adv 2:e1600584. 10.1126/sciadv.1600584

6. Kennedy BK, Berger SL, Brunet A, Campisi J, Cuervo AM, Epel ES, Franceschi C, Lithgow GJ, Morimoto RI, Pessin JE, Rando TA, Richardson A, Schadt EE, Wyss-Coray T, Sierra F (2014) Aging: a common driver of chronic diseases and a target for novel interventions. Cell 159:709–713. 10.1016/j.cell.2014.10.039

7. Wu Z, Qu J, Zhang W, Liu G-H (2024) Stress, epigenetics, and aging: Unraveling the intricate crosstalk. Molecular Cell 84:34–54. 10.1016/j.molcel.2023.10.006

8. Kane AE, Sinclair DA (2019) Epigenetic changes during aging and their reprogramming potential. Critical Reviews in Biochemistry and Molecular Biology

9. Moskalev A, Stambler I, Caruso C (2020) Innate and Adaptive Immunity in Aging and Longevity: The Foundation of Resilience. Aging Dis 11:1363–1373. 10.14336/AD.2020.0603

10. Solana R, Tarazona R, Gayoso I, Lesur O, Dupuis G, Fulop T (2012) Innate immunosenescence: effect of aging on cells and receptors of the innate immune system in humans. Semin Immunol 24:331–341. 10.1016/j.smim.2012.04.008

11. Nikolich-Žugich J (2018) The twilight of immunity: emerging concepts in aging of the immune system. Nat Immunol 19:10–19. 10.1038/s41590-017-0006-x

12. Fulop T, Larbi A, Pawelec G, Khalil A, Cohen AA, Hirokawa K, Witkowski JM, Franceschi C (2023) Immunology of Aging: the Birth of Inflammaging. Clinical Reviews in Allergy & Immunology 64:109. 10.1007/s12016-021-08899-6

13. Rea IM, Gibson DS, McGilligan V, McNerlan SE, Alexander HD, Ross OA (2018) Age and Age-Related Diseases: Role of Inflammation Triggers and Cytokines. Front Immunol 9:586. 10.3389/fimmu.2018.00586

14. Franceschi C, Bonafè M, Valensin S, Olivieri F, De Luca M, Ottaviani E, De Benedictis G (2000) Inflamm-aging. An evolutionary perspective on immunosenescence. Ann N Y Acad Sci 908:244–254. 10.1111/j.1749-6632.2000.tb06651.x

15. Bacos K, Gillberg L, Volkov P, Olsson AH, Hansen T, Pedersen O, Gjesing AP, Eiberg H, Tuomi T, Almgren P, Groop L, Eliasson L, Vaag A, Dayeh T, Ling C (2016) Blood-based biomarkers of age-associated epigenetic changes in human islets associate with insulin secretion and diabetes. Nat Commun 7:11089. 10.1038/ncomms11089

16. Rönn T, Volkov P, Gillberg L, Kokosar M, Perfilyev A, Jacobsen AL, Jørgensen SW, Brøns C, Jansson P-A, Eriksson K-F, Pedersen O, Hansen T, Groop L, Stener-Victorin E, Vaag A, Nilsson E, Ling C (2015) Impact of age, BMI and HbA1c levels on the genome-wide DNA methylation and mRNA expression patterns in human adipose tissue and identification of epigenetic biomarkers in blood. Human Molecular Genetics 24:3792–3813. 10.1093/hmg/ddv124

17. Bacalini MG, Deelen J, Pirazzini C, De Cecco M, Giuliani C, Lanzarini C, Ravaioli F, Marasco E, van Heemst D, Suchiman HED, Slieker R, Giampieri E, Recchioni R, Mercheselli F, Salvioli S, Vitale G, Olivieri F, Spijkerman AMW, Dollé MET, Sedivy JM, Castellani G, Franceschi C, Slagboom PE, Garagnani P (2017) Systemic Age-Associated DNA Hypermethylation of ELOVL2 Gene: In Vivo and In Vitro Evidences of a Cell Replication Process. The Journals of Gerontology: Series A 72:1015–1023. 10.1093/gerona/glw185

18. Slieker RC, Relton CL, Gaunt TR, Slagboom PE, Heijmans BT (2018) Age-related DNA methylation changes are tissue-specific with ELOVL2 promoter methylation as exception. Epigenetics & Chromatin 11:25. 10.1186/s13072-018-0191-3

19. Nardini C, Moreau J-F, Gensous N, Ravaioli F, Garagnani P, Bacalini MG (2018) The epigenetics of inflammaging: The contribution of age-related heterochromatin loss and locus-specific remodelling and the modulation by environmental stimuli. Seminars in Immunology 40:49–60. 10.1016/j.smim.2018.10.009

20. Agrawal A, Tay J, Yang G-E, Agrawal S, Gupta S (2010) Age-associated epigenetic modifications in human DNA increase its immunogenicity. Aging 2:93–100. 10.18632/aging.100121

21. Jylhävä J, Pedersen NL, Hägg S (2017) Biological Age Predictors. eBioMedicine 21:29–36. 10.1016/j.ebiom.2017.03.046

22. Horvath S (2013) DNA methylation age of human tissues and cell types. Genome Biol 14:R115. 10.1186/gb-2013-14-10-r115

23. Hannum G, Guinney J, Zhao L, Zhang L, Hughes G, Sadda S, Klotzle B, Bibikova M, Fan J-B, Gao Y, Deconde R, Chen M, Rajapakse I, Friend S, Ideker T, Zhang K (2013) Genome-wide Methylation Profiles Reveal Quantitative Views of Human Aging Rates. Molecular Cell 49:359–367. 10.1016/j.molcel.2012.10.016

24. Levine ME, Lu AT, Quach A, Chen BH, Assimes TL, Bandinelli S, Hou L, Baccarelli AA, Stewart JD, Li Y, Whitsel EA, Wilson JG, Reiner AP, Aviv A, Lohman K, Liu Y, Ferrucci L, Horvath S (2018) An epigenetic biomarker of aging for lifespan and healthspan. Aging 10:573–591. 10.18632/aging.101414

25. Lu AT, Quach A, Wilson JG, Reiner AP, Aviv A, Raj K, Hou L, Baccarelli AA, Li Y, Stewart JD, Whitsel EA, Assimes TL, Ferrucci L, Horvath S (2019) DNA methylation GrimAge strongly predicts lifespan and healthspan. Aging 11:303–327. 10.18632/aging.101684

26. Belsky DW, Caspi A, Arseneault L, Baccarelli A, Corcoran DL, Gao X, Hannon E, Harrington HL, Rasmussen LJ, Houts R, Huffman K, Kraus WE, Kwon D, Mill J, Pieper CF, Prinz JA, Poulton R, Schwartz J, Sugden K, Vokonas P, Williams BS, Moffitt TE (2020) Quantification of the pace of biological aging in humans through a blood test, the DunedinPoAm DNA methylation algorithm. Elife 9:e54870. 10.7554/eLife.54870

27. Belsky DW, Caspi A, Corcoran DL, Sugden K, Poulton R, Arseneault L, Baccarelli A, Chamarti K, Gao X, Hannon E, Harrington HL, Houts R, Kothari M, Kwon D, Mill J, Schwartz J, Vokonas P, Wang C, Williams BS, Moffitt TE (2022) DunedinPACE, a DNA methylation biomarker of the pace of aging. eLife 11:e73420. 10.7554/eLife.73420

28. Ying K, Liu H, Tarkhov AE, Sadler MC, Lu AT, Moqri M, Horvath S, Kutalik Z, Shen X, Gladyshev VN (2024) Causality-enriched epigenetic age uncouples damage and adaptation. Nat Aging 4:231–246. 10.1038/s43587-023-00557-0

29. Oblak L, van der Zaag J, Higgins-Chen AT, Levine ME, Boks MP (2021) A systematic review of biological, social and environmental factors associated with epigenetic clock acceleration. Ageing Research Reviews 69:101348. 10.1016/j.arr.2021.101348

30. Fransquet PD, Wrigglesworth J, Woods RL, Ernst ME, Ryan J (2019) The epigenetic clock as a predictor of disease and mortality risk: a systematic review and meta-analysis. Clin Epigenetics 11:62. 10.1186/s13148-019-0656-7

31. Cao X, Li W, Wang T, Ran D, Davalos V, Planas-Serra L, Pujol A, Esteller M, Wang X, Yu H (2022) Accelerated biological aging in COVID-19 patients. Nat Commun 13:2135. 10.1038/s41467-022-29801-8

32. Sayed N, Huang Y, Nguyen K, Krejciova-Rajaniemi Z, Grawe AP, Gao T, Tibshirani R, Hastie T, Alpert A, Cui L, Kuznetsova T, Rosenberg-Hasson Y, Ostan R, Monti D, Lehallier B, Shen-Orr SS, Maecker HT, Dekker CL, Wyss-Coray T, Franceschi C, Jojic V, Haddad F, Montoya JG, Wu JC, Davis MM, Furman D (2021) An inflammatory aging clock (iAge) based on deep learning tracks multimorbidity, immunosenescence, frailty and cardiovascular aging. Nat Aging 1:598–615. 10.1038/s43587-021-00082-y

33. Kalyakulina A, Yusipov I, Kondakova E, Bacalini MG, Franceschi C, Vedunova M, Ivanchenko M (2023) Small immunological clocks identified by deep learning and gradient boosting. Front Immunol 14:. 10.3389/fimmu.2023.1177611

34. Murabito JM, Zhao Q, Larson MG, Rong J, Lin H, Benjamin EJ, Levy D, Lunetta KL (2018) Measures of Biologic Age in a Community Sample Predict Mortality and Age-Related Disease: The Framingham Offspring Study. The Journals of Gerontology: Series A 73:757–762. 10.1093/gerona/glx144

35. Alpert A, Pickman Y, Leipold M, Rosenberg-Hasson Y, Ji X, Gaujoux R, Rabani H, Starosvetsky E, Kveler K, Schaffert S, Furman D, Caspi O, Rosenschein U, Khatri P, Dekker CL, Maecker HT, Davis MM, Shen-Orr SS (2019) A clinically meaningful metric of immune age derived from high-dimensional longitudinal monitoring. Nat Med 25:487–495. 10.1038/s41591-019-0381-y

36. Yusipov I, Kondakova E, Kalyakulina A, Krivonosov M, Lobanova N, Bacalini MG, Franceschi C, Vedunova M, Ivanchenko M (2022) Accelerated epigenetic aging and inflammatory/immunological profile (ipAGE) in patients with chronic kidney disease. Geroscience 44:817–834. 10.1007/s11357-022-00540-4

37. Markov NT, Lindbergh CA, Staffaroni AM, Perez K, Stevens M, Nguyen K, Murad NF, Fonseca C, Campisi J, Kramer J, Furman D (2022) Age-related brain atrophy is not a homogenous process: Different functional brain networks associate differentially with aging and blood factors. Proceedings of the National Academy of Sciences 119:e2207181119. 10.1073/pnas.2207181119

38. Crimmins EM, Klopack ET, Kim JK (2024) Generations of epigenetic clocks and their links to socioeconomic status in the Health and Retirement Study. Epigenomics 16:1031–1042. 10.1080/17501911.2024.2373682

39. Yusipov I, Kalyakulina A, Trukhanov A, Franceschi C, Ivanchenko M (2024) Map of epigenetic age acceleration: A worldwide analysis. Ageing Research Reviews 100:102418. 10.1016/j.arr.2024.102418

40. Horvath S, Raj K (2018) DNA methylation-based biomarkers and the epigenetic clock theory of ageing. Nat Rev Genet 19:371–384. 10.1038/s41576-018-0004-3

41. Gadd DA, Hillary RF, McCartney DL, Zaghlool SB, Stevenson AJ, Cheng Y, Fawns-Ritchie C, Nangle C, Campbell A, Flaig R, Harris SE, Walker RM, Shi L, Tucker-Drob EM, Gieger C, Peters A, Waldenberger M, Graumann J, McRae AF, Deary IJ, Porteous DJ, Hayward C, Visscher PM, Cox SR, Evans KL, McIntosh AM, Suhre K, Marioni RE (2022) Epigenetic scores for the circulating proteome as tools for disease prediction. Elife 11:e71802. 10.7554/eLife.71802

42. Bernabeu E, McCartney DL, Gadd DA, Hillary RF, Lu AT, Murphy L, Wrobel N, Campbell A, Harris SE, Liewald D, Hayward C, Sudlow C, Cox SR, Evans KL, Horvath S, McIntosh AM, Robinson MR, Vallejos CA, Marioni RE (2023) Refining epigenetic prediction of chronological and biological age. Genome Med 15:12. 10.1186/s13073-023-01161-y

43. Kalyakulina A, Yusipov I (2024) Web-interface for calculating EpImAge. In: EpImAge. https://huggingface.co/spaces/UNNAILab/EpImAge. Accessed 16 Oct 2024

44. Pidsley R, Zotenko E, Peters TJ, Lawrence MG, Risbridger GP, Molloy P, Van Djik S, Muhlhausler B, Stirzaker C, Clark SJ (2016) Critical evaluation of the Illumina MethylationEPIC BeadChip microarray for whole-genome DNA methylation profiling. Genome Biol 17:208. 10.1186/s13059-016-1066-1

45. Tian Y, Morris TJ, Webster AP, Yang Z, Beck S, Feber A, Teschendorff AE (2017) ChAMP: updated methylation analysis pipeline for Illumina BeadChips. Bioinformatics 33:3982–3984. 10.1093/bioinformatics/btx513

46. Aryee MJ, Jaffe AE, Corrada-Bravo H, Ladd-Acosta C, Feinberg AP, Hansen KD, Irizarry RA (2014) Minfi: a flexible and comprehensive Bioconductor package for the analysis of Infinium DNA methylation microarrays. Bioinformatics 30:1363–1369. 10.1093/bioinformatics/btu049

47. Zhu Y, Davis S, Stephens R, Meltzer PS, Chen Y (2008) GEOmetadb: powerful alternative search engine for the Gene Expression Omnibus. Bioinformatics 24:2798– 2800. 10.1093/bioinformatics/btn520

48. Davis S, Meltzer PS (2007) GEOquery: a bridge between the Gene Expression Omnibus (GEO) and BioConductor. Bioinformatics 23:1846–1847. 10.1093/bioinformatics/btm254

49. Barrett T, Wilhite SE, Ledoux P, Evangelista C, Kim IF, Tomashevsky M, Marshall KA, Phillippy KH, Sherman PM, Holko M, Yefanov A, Lee H, Zhang N, Robertson CL, Serova N, Davis S, Soboleva A (2013) NCBI GEO: archive for functional genomics data sets—update. Nucleic Acids Research 41:D991–D995. 10.1093/nar/gks1193

50. Gumienny R, van Heeringen S, Bismeijer T, Ramdhani H (2021) GEOparse: Python library to access Gene Expression Omnibus Database (GEO)

51. Ding C, Peng H (2003) Minimum redundancy feature selection from microarray gene expression data. In: Computational Systems Bioinformatics. CSB2003. Proceedings of the 2003 IEEE Bioinformatics Conference. CSB2003. pp 523–528

52. Chen J, Liao K, Wan Y, Chen DZ, Wu J (2022) DANets: Deep Abstract Networks for Tabular Data Classification and Regression. AAAI 36:3930–3938. 10.1609/aaai.v36i4.20309

53. Gorishniy Y, Rubachev I, Khrulkov V, Babenko A (2021) Revisiting Deep Learning Models for Tabular Data. In: Advances in Neural Information Processing Systems. Curran Associates, Inc., pp 18932–18943

54. Joseph M, Raj H (2024) GANDALF: Gated Adaptive Network for Deep Automated Learning of Features

55. Lundberg SM, Lee S-I (2017) A unified approach to interpreting model predictions. In: Proceedings of the 31st International Conference on Neural Information Processing Systems. Curran Associates Inc., Red Hook, NY, USA, pp 4768–4777

56. Mann HB, Whitney DR (1947) On a Test of Whether one of Two Random Variables is Stochastically Larger than the Other. The Annals of Mathematical Statistics 18:50–60. 10.1214/aoms/1177730491

57. Benjamini Y, Hochberg Y (1995) Controlling the False Discovery Rate: A Practical and Powerful Approach to Multiple Testing. Journal of the Royal Statistical Society: Series B (Methodological) 57:289–300. 10.1111/j.2517-6161.1995.tb02031.x

58. de Lima Camillo LP (2024) pyaging: a Python-based compendium of GPU-optimized aging clocks. Bioinformatics 40:btae200. 10.1093/bioinformatics/btae200

59. Lin Q, Weidner CI, Costa IG, Marioni RE, Ferreira MRP, Deary IJ, Wagner W (2016) DNA methylation levels at individual age-associated CpG sites can be indicative for life expectancy. Aging 8:394–401. 10.18632/aging.100908

60. Yang Z, Wong A, Kuh D, Paul DS, Rakyan VK, Leslie RD, Zheng SC, Widschwendter M, Beck S, Teschendorff AE (2016) Correlation of an epigenetic mitotic clock with cancer risk. Genome Biology 17:205. 10.1186/s13059-016-1064-3

61. Zhang Y, Wilson R, Heiss J, Breitling LP, Saum K-U, Schöttker B, Holleczek B, Waldenberger M, Peters A, Brenner H (2017) DNA methylation signatures in peripheral blood strongly predict all-cause mortality. Nat Commun 8:14617. 10.1038/ncomms14617

62. Horvath S, Oshima J, Martin GM, Lu AT, Quach A, Cohen H, Felton S, Matsuyama M, Lowe D, Kabacik S, Wilson JG, Reiner AP, Maierhofer A, Flunkert J, Aviv A, Hou L, Baccarelli AA, Li Y, Stewart JD, Whitsel EA, Ferrucci L, Matsuyama S, Raj K (2018) Epigenetic clock for skin and blood cells applied to Hutchinson Gilford Progeria Syndrome and *ex vivo* studies. Aging 10:1758–1775. 10.18632/aging.101508

63. Lu AT, Seeboth A, Tsai P-C, Sun D, Quach A, Reiner AP, Kooperberg C, Ferrucci L, Hou L, Baccarelli AA, Li Y, Harris SE, Corley J, Taylor A, Deary IJ, Stewart JD, Whitsel EA, Assimes TL, Chen W, Li S, Mangino M, Bell JT, Wilson JG, Aviv A, Marioni RE, Raj K, Horvath S (2019) DNA methylation-based estimator of telomere length. Aging 11:5895–5923. 10.18632/aging.102173

64. Zhang Q, Vallerga CL, Walker RM, Lin T, Henders AK, Montgomery GW, He J, Fan D, Fowdar J, Kennedy M, Pitcher T, Pearson J, Halliday G, Kwok JB, Hickie I, Lewis S, Anderson T, Silburn PA, Mellick GD, Harris SE, Redmond P, Murray AD, Porteous DJ, Haley CS, Evans KL, McIntosh AM, Yang J, Gratten J, Marioni RE, Wray NR, Deary IJ, McRae AF, Visscher PM (2019) Improved precision of epigenetic clock estimates across tissues and its implication for biological ageing. Genome Medicine 11:54. 10.1186/s13073-019-0667-1

65. Han Y, Franzen J, Stiehl T, Gobs M, Kuo C-C, Nikolić M, Hapala J, Koop BE, Strathmann K, Ritz-Timme S, Wagner W (2020) New targeted approaches for epigenetic age predictions. BMC Biology 18:71. 10.1186/s12915-020-00807-2

66. de Lima Camillo LP, Lapierre LR, Singh R (2022) A pan-tissue DNA-methylation epigenetic clock based on deep learning. npj Aging 8:1–15. 10.1038/s41514-022-00085-y

67. Higgins-Chen AT, Thrush KL, Wang Y, Minteer CJ, Kuo P-L, Wang M, Niimi P, Sturm G, Lin J, Moore AZ, Bandinelli S, Vinkers CH, Vermetten E, Rutten BPF, Geuze E, Okhuijsen-Pfeifer C, van der Horst MZ, Schreiter S, Gutwinski S, Luykx JJ, Picard M, Ferrucci L, Crimmins EM, Boks MP, Hägg S, Hu-Seliger TT, Levine ME (2022) A computational solution for bolstering reliability of epigenetic clocks: implications for clinical trials and longitudinal tracking. Nat Aging 2:644–661. 10.1038/s43587-022-00248-2

68. Lu AT, Binder AM, Zhang J, Yan Q, Reiner AP, Cox SR, Corley J, Harris SE, Kuo P- L, Moore AZ, Bandinelli S, Stewart JD, Wang C, Hamlat EJ, Epel ES, Schwartz JD, Whitsel EA, Correa A, Ferrucci L, Marioni RE, Horvath S (2022) DNA methylation GrimAge version 2. Aging 14:9484–9549. 10.18632/aging.204434

69. McGreevy KM, Radak Z, Torma F, Jokai M, Lu AT, Belsky DW, Binder A, Marioni RE, Ferrucci L, Pośpiech E, Branicki W, Ossowski A, Sitek A, Spólnicka M, Raffield LM, Reiner AP, Cox S, Kobor M, Corcoran DL, Horvath S (2023) DNAmFitAge: biological age indicator incorporating physical fitness. Aging (Albany NY) 15:3904– 3938. 10.18632/aging.204538

70. Dec E, Clement J, Cheng K, Church GM, Fossel MB, Rehkopf DH, Rosero-Bixby L, Kobor MS, Lin DTS, Lu AT, Fei Z, Guo W, Chew YC, Yang X, Putra SED, Reiner AP, Correa A, Vilalta A, Pirazzini C, Passarino G, Monti D, Arosio B, Garagnani P, Franceschi C, Horvath S (2023) Centenarian clocks: epigenetic clocks for validating claims of exceptional longevity. GeroScience 45:1817–1835. 10.1007/s11357-023-00731-7

71. Tong H, Dwaraka VB, Chen Q, Luo Q, Lasky-Su JA, Smith R, Teschendorff AE (2024) Quantifying the stochastic component of epigenetic aging. Nat Aging 4:886–901. 10.1038/s43587-024-00600-8

72. Zhu T, Tong H, Du Z, Beck S, Teschendorff AE (2024) An improved epigenetic counter to track mitotic age in normal and precancerous tissues. Nat Commun 15:4211. 10.1038/s41467-024-48649-8

73. Ndhlovu LC, Bendall ML, Dwaraka V, Pang APS, Dopkins N, Carreras N, Smith R, Nixon DF, Corley MJ (2024) Retro-age: A unique epigenetic biomarker of aging captured by DNA methylation states of retroelements. Aging Cell e14288. 10.1111/acel.14288

74. Tomusiak A, Floro A, Tiwari R, Riley R, Matsui H, Andrews N, Kasler HG, Verdin E (2024) Development of an epigenetic clock resistant to changes in immune cell composition. Commun Biol 7:1–13. 10.1038/s42003-024-06609-4

75. Chen EY, Tan CM, Kou Y, Duan Q, Wang Z, Meirelles GV, Clark NR, Ma’ayan A (2013) Enrichr: interactive and collaborative HTML5 gene list enrichment analysis tool. BMC Bioinformatics 14:128. 10.1186/1471-2105-14-128

76. Kuleshov MV, Jones MR, Rouillard AD, Fernandez NF, Duan Q, Wang Z, Koplev S, Jenkins SL, Jagodnik KM, Lachmann A, McDermott MG, Monteiro CD, Gundersen GW, Ma’ayan A (2016) Enrichr: a comprehensive gene set enrichment analysis web server 2016 update. Nucleic Acids Res 44:W90–97. 10.1093/nar/gkw377

77. Xie Z, Bailey A, Kuleshov MV, Clarke DJB, Evangelista JE, Jenkins SL, Lachmann A, Wojciechowicz ML, Kropiwnicki E, Jagodnik KM, Jeon M, Ma’ayan A (2021) Gene Set Knowledge Discovery with Enrichr. Current Protocols 1:e90. 10.1002/cpz1.90

78. Abid A, Abdalla A, Abid A, Khan D, Alfozan A, Zou J (2019) Gradio: Hassle-Free Sharing and Testing of ML Models in the Wild

79. Kalyakulina A, Yusipov I, Kondakova E, Bacalini MG, Giuliani C, Sivtseva T, Semenov S, Ksenofontov A, Nikolaeva M, Khusnutdinova E, Zakharova R, Vedunova M, Franceschi C, Ivanchenko M (2023) Epigenetics of the far northern Yakutian population. Clinical Epigenetics 15:189. 10.1186/s13148-023-01600-y

80. Ding C, Peng H (2005) Minimum redundancy feature selection from microarray gene expression data. J Bioinform Comput Biol 3:185–205. 10.1142/s0219720005001004

81. Gene Ontology Consortium (2021) The Gene Ontology resource: enriching a GOld mine. Nucleic Acids Res 49:D325–D334. 10.1093/nar/gkaa1113

82. Kanehisa M, Goto S (2000) KEGG: kyoto encyclopedia of genes and genomes. Nucleic Acids Res 28:27–30. 10.1093/nar/28.1.27

83. Zeitz MJ, Smyth JW (2023) Gap Junctions and Ageing. Subcell Biochem 102:113–137. 10.1007/978-3-031-21410-3_6

84. Dong D, Xie W, Liu M (2020) Alteration of cell junctions during viral infection. Thorac Cancer 11:519–525. 10.1111/1759-7714.13344

85. Ashcroft GS, Horan MA, Ferguson MW (1998) Aging alters the inflammatory and endothelial cell adhesion molecule profiles during human cutaneous wound healing. Lab Invest 78:47–58

86. Arnesen SM, Lawson MA (2006) Age-related changes in focal adhesions lead to altered cell behavior in tendon fibroblasts. Mech Ageing Dev 127:726–732. 10.1016/j.mad.2006.05.003

87. Harjunpää H, Llort Asens M, Guenther C, Fagerholm SC (2019) Cell Adhesion Molecules and Their Roles and Regulation in the Immune and Tumor Microenvironment. Front Immunol 10:1078. 10.3389/fimmu.2019.01078

88. Salvadores N, Sanhueza M, Manque P, Court FA (2017) Axonal Degeneration during Aging and Its Functional Role in Neurodegenerative Disorders. Front Neurosci 11:451. 10.3389/fnins.2017.00451

89. Azpurua J, Eaton BA (2015) Neuronal epigenetics and the aging synapse. Front Cell Neurosci 9:. 10.3389/fncel.2015.00208

90. Galloway DA, Phillips AEM, Owen DRJ, Moore CS (2019) Phagocytosis in the Brain: Homeostasis and Disease. Front Immunol 10:790. 10.3389/fimmu.2019.00790

91. Li W (2013) Phagocyte dysfunction, tissue aging and degeneration. Ageing Res Rev 12:10.1016/j.arr.2013.05.006. 10.1016/j.arr.2013.05.006

92. Frobel J, Hemeda H, Lenz M, Abagnale G, Joussen S, Denecke B, Šarić T, Zenke M, Wagner W (2014) Epigenetic Rejuvenation of Mesenchymal Stromal Cells Derived from Induced Pluripotent Stem Cells. Stem Cell Reports 3:414–422. 10.1016/j.stemcr.2014.07.003

93. Lodde V, Floris M, Munk R, Martindale JL, Piredda D, Napodano CMP, Cucca F, Uzzau S, Abdelmohsen K, Gorospe M, Noh JH, Idda ML (2022) Systematic identification of NF90 target RNAs by iCLIP analysis. Sci Rep 12:364. 10.1038/s41598-021-04101-1

94. Păun O, Tan YX, Patel H, Strohbuecker S, Ghanate A, Cobolli-Gigli C, Llorian Sopena M, Gerontogianni L, Goldstone R, Ang S-L, Guillemot F, Dias C (2023) Pioneer factor ASCL1 cooperates with the mSWI/SNF complex at distal regulatory elements to regulate human neural differentiation. Genes Dev 37:218–242. 10.1101/gad.350269.122

95. Marttila S, Chatsirisupachai K, Palmer D, de Magalhães JP (2020) Ageing-associated changes in the expression of lncRNAs in human tissues reflect a transcriptional modulation in ageing pathways. Mech Ageing Dev 185:111177. 10.1016/j.mad.2019.111177

96. Di Sabatino A, Calarota SA, Vidali F, MacDonald TT, Corazza GR (2011) Role of IL-15 in immune-mediated and infectious diseases. Cytokine & Growth Factor Reviews 22:19–33. 10.1016/j.cytogfr.2010.09.003

97. Loughland JR, Woodberry T, Oyong D, Piera KA, Amante FH, Barber BE, Grigg MJ, William T, Engwerda CR, Anstey NM, McCarthy JS, Boyle MJ, Minigo G (2021) Reduced circulating dendritic cells in acute Plasmodium knowlesi and Plasmodium falciparum malaria despite elevated plasma Flt3 ligand levels. Malaria Journal 20:97. 10.1186/s12936-021-03642-0

98. Mahittikorn A, Kwankaew P, Rattaprasert P, Kotepui KU, Masangkay FR, Kotepui M (2022) Elevation of serum interleukin-1β levels as a potential indicator for malarial infection and severe malaria: a meta-analysis. Malaria Journal 21:308. 10.1186/s12936-022-04325-0

99. Liu M, Guo S, Hibbert JM, Jain V, Singh N, Wilson NO, Stiles JK (2011) CXCL10/IP-10 in infectious diseases pathogenesis and potential therapeutic implications. Cytokine & Growth Factor Reviews 22:121. 10.1016/j.cytogfr.2011.06.001

100. Quartuccio L, Fabris M, Sonaglia A, Peghin M, Domenis R, Cifù A, Curcio F, Tascini C (2021) Interleukin 6, soluble interleukin 2 receptor alpha (CD25), monocyte colony-stimulating factor, and hepatocyte growth factor linked with systemic hyperinflammation, innate immunity hyperactivation, and organ damage in COVID-19 pneumonia. Cytokine 140:155438. 10.1016/j.cyto.2021.155438

101. Korobova ZR, Arsentieva NA, Liubimova NE, Dedkov VG, Gladkikh AS, Sharova AA, Chernykh EI, Kashchenko VA, Ratnikov VA, Gorelov VP, Stanevich OV, Kulikov AN, Pevtsov DE, Totolian AA (2022) A Comparative Study of the Plasma Chemokine Profile in COVID-19 Patients Infected with Different SARS-CoV-2 Variants. Int J Mol Sci 23:9058. 10.3390/ijms23169058

102. D’Rozario R, Raychaudhuri D, Bandopadhyay P, Sarif J, Mehta P, Liu CSC, Sinha BP, Roy J, Bhaduri R, Das M, Bandyopadhyay S, Paul SR, Chatterjee S, Pandey R, Ray Y, Ganguly D (2023) Circulating Interleukin-8 Dynamics Parallels Disease Course and Is Linked to Clinical Outcomes in Severe COVID-19. Viruses 15:549. 10.3390/v15020549

103. Mandel M, Harari G, Gurevich M, Achiron A (2020) Cytokine prediction of mortality in COVID19 patients. Cytokine 134:155190. 10.1016/j.cyto.2020.155190

104. Pius-Sadowska E, Niedźwiedź A, Kulig P, Baumert B, Sobuś A, Rogińska D, Łuczkowska K, Ulańczyk Z, Wnęk S, Karolak I, Paczkowska E, Kotfis K, Kawa M, Stecewicz I, Zawodny P, Machaliński B (2022) CXCL8, CCL2, and CMV Seropositivity as New Prognostic Factors for a Severe COVID-19 Course. Int J Mol Sci 23:11338. 10.3390/ijms231911338

105. Khalil BA, Elemam NM, Maghazachi AA (2021) Chemokines and chemokine receptors during COVID-19 infection. Computational and Structural Biotechnology Journal 19:976–988. 10.1016/j.csbj.2021.01.034

106. Bekbossynova M, Tauekelova A, Sailybayeva A, Kozhakhmetov S, Mussabay K, Chulenbayeva L, Kossumov A, Khassenbekova Z, Vinogradova E, Kushugulova A (2023) Unraveling Acute and Post-COVID Cytokine Patterns to Anticipate Future Challenges. J Clin Med 12:5224. 10.3390/jcm12165224

107. Pan T, Gallo ME, Donald KA, Webb K, Bath KG (2024) Elevated risk for psychiatric outcomes in pediatric patients with Multisystem Inflammatory Syndrome (MIS-C): A review of neuroinflammatory and psychosocial stressors. Brain, Behavior, & Immunity - Health 38:100760. 10.1016/j.bbih.2024.100760

108. Kameda M, Otsuka M, Chiba H, Kuronuma K, Hasegawa T, Takahashi H, Takahashi H (2020) CXCL9, CXCL10, and CXCL11; biomarkers of pulmonary inflammation associated with autoimmunity in patients with collagen vascular diseases–associated interstitial lung disease and interstitial pneumonia with autoimmune features. PLoS One 15:e0241719. 10.1371/journal.pone.0241719

109. Xu J, Zhong S, Liu J, Li L, Li Y, Wu X, Li Z, Deng P, Zhang J, Zhong N, Ding Y, Jiang Y (2005) Detection of Severe Acute Respiratory Syndrome Coronavirus in the Brain: Potential Role of the Chemokine Mig in Pathogenesis. Clinical Infectious Diseases 41:1089–1096. 10.1086/444461

110. Stefano AD, Coccini T, Roda E, Signorini C, Balbi B, Brunetti G, Ceriana P (2018) Blood MCP-1 levels are increased in chronic obstructive pulmonary disease patients with prevalent emphysema. International Journal of Chronic Obstructive Pulmonary Disease 13:1691. 10.2147/COPD.S159915

111. Reynolds D, Vazquez Guillamet C, Day A, Borcherding N, Vazquez Guillamet R, Choreño-Parra JA, House SL, O’Halloran JA, Zúñiga J, Ellebedy AH, Byers DE, Mudd PA (2021) Comprehensive Immunologic Evaluation of Bronchoalveolar Lavage Samples from Human Patients with Moderate and Severe Seasonal Influenza and Severe COVID-19. The Journal of Immunology 207:1229–1238. 10.4049/jimmunol.2100294

112. Rincon M, Irvin CG (2012) Role of IL-6 in Asthma and Other Inflammatory Pulmonary Diseases. International Journal of Biological Sciences 8:1281. 10.7150/ijbs.4874

113. Moledina DG, Obeid W, Smith RN, Rosales I, Sise ME, Moeckel G, Kashgarian M, Kuperman M, Campbell KN, Lefferts S, Meliambro K, Bitzer M, Perazella MA, Luciano RL, Pober JS, Cantley LG, Colvin RB, Wilson FP, Parikh CR Identification and validation of urinary CXCL9 as a biomarker for diagnosis of acute interstitial nephritis. J Clin Invest 133:e168950. 10.1172/JCI168950

114. Araújo LS, Torquato BGS, da Silva CA, dos Reis Monteiro MLG, dos Santos Martins ALM, da Silva MV, dos Reis MA, Machado JR (2020) Renal expression of cytokines and chemokines in diabetic nephropathy. BMC Nephrol 21:308. 10.1186/s12882-020-01960-0

115. Roy MS, Janal MN, Crosby J, Donnelly R (2015) Markers of endothelial dysfunction and inflammation predict progression of diabetic nephropathy in African Americans with type 1 diabetes. Kidney International 87:427–433. 10.1038/ki.2014.212

116. de Oliveira Junior WV, Silva APF, de Figueiredo RC, Gomes KB, Simões e Silva AC, Dusse LMS, Rios DRA (2020) Association between dyslipidemia and CCL2 in patients undergoing hemodialysis. Cytokine 125:154858. 10.1016/j.cyto.2019.154858

117. Badeński A, Badeńska M, Świętochowska E, Janek A, Gliwińska A, Morawiec-Knysak A, Szczepańska M (2023) Assessment of Interleukin-15 (IL-15) Concentration in Children with Idiopathic Nephrotic Syndrome. Int J Mol Sci 24:6993. 10.3390/ijms24086993

118. Lebherz-Eichinger D, Klaus DA, Reiter T, Hörl WH, Haas M, Ankersmit HJ, Krenn CG, Roth GA (2014) Increased chemokine excretion in patients suffering from chronic kidney disease. Translational Research 164:433–443.e2. 10.1016/j.trsl.2014.07.004

119. Fallahi P, Ferrari SM, Ragusa F, Ruffilli I, Elia G, Paparo SR, Antonelli A (2020) Th1 Chemokines in Autoimmune Endocrine Disorders. The Journal of Clinical Endocrinology & Metabolism 105:1046–1060. 10.1210/clinem/dgz289

120. Ullah A, Ud Din A, Ding W, Shi Z, Pervaz S, Shen B (2023) A narrative review: CXC chemokines influence immune surveillance in obesity and obesity-related diseases: Type 2 diabetes and nonalcoholic fatty liver disease. Rev Endocr Metab Disord 24:611–631. 10.1007/s11154-023-09800-w

121. Bastard J-P, Jardel C, Bruckert E, Blondy P, Capeau J, Laville M, Vidal H, Hainque B (2000) Elevated Levels of Interleukin 6 Are Reduced in Serum and Subcutaneous Adipose Tissue of Obese Women after Weight Loss. The Journal of Clinical Endocrinology & Metabolism 85:3338–3342. 10.1210/jcem.85.9.6839

122. Shen Y, Ou Ji, Liu M, Shi L, Li Y, Xiao L, Dong H, Zhang F, Xia K, Zhao J (2016) Altered plasma levels of chemokines in autism and their association with social behaviors. Psychiatry Research 244:300–305. 10.1016/j.psychres.2016.07.057

123. Zhou X, Tian B, Han H-B (2021) Serum interleukin-6 in schizophrenia: A system review and meta-analysis. Cytokine 141:155441. 10.1016/j.cyto.2021.155441

124. Roohi E, Jaafari N, Hashemian F (2021) On inflammatory hypothesis of depression: what is the role of IL-6 in the middle of the chaos? Journal of Neuroinflammation 18:45. 10.1186/s12974-021-02100-7

125. Ridker PM, Rane M (2021) Interleukin-6 Signaling and Anti-Interleukin-6 Therapeutics in Cardiovascular Disease. Circulation Research 128:1728–1746. 10.1161/CIRCRESAHA.121.319077

126. Cainzos-Achirica M, Enjuanes C, Greenland P, McEvoy JW, Cushman M, Dardari Z, Nasir K, Budoff MJ, Al-Mallah MH, Yeboah J, Miedema MD, Blumenthal RS, Comin-Colet J, Blaha MJ (2018) The prognostic value of interleukin 6 in multiple chronic diseases and all-cause death: The Multi-Ethnic Study of Atherosclerosis (MESA). Atherosclerosis 278:217–225. 10.1016/j.atherosclerosis.2018.09.034

127. Eindor A, Tsai K, Jacobson K (2024) P125 MIG (CXCL9) and IL22 are key biomarkers that discriminates between paediatric IBD patients and non-IBD patients in a novel biomarker model. Journal of Crohn’s and Colitis 18:i418. 10.1093/ecco-jcc/jjad212.0255

128. Jarlborg M, Gabay C (2022) Systemic effects of IL-6 blockade in rheumatoid arthritis beyond the joints. Cytokine 149:155742. 10.1016/j.cyto.2021.155742

129. Schoepf IC, Esteban-Cantos A, Thorball CW, Rodés B, Reiss P, Rodríguez-Centeno J, Riebensahm C, Braun DL, Marzolini C, Seneghini M, Bernasconi E, Cavassini M, Buvelot H, Thurnheer MC, Kouyos RD, Fellay J, Günthard HF, Arribas JR, Ledergerber B, Tarr PE (2023) Epigenetic ageing accelerates before antiretroviral therapy and decelerates after viral suppression in people with HIV in Switzerland: a longitudinal study over 17 years. The Lancet Healthy Longevity 4:e211–e218. 10.1016/S2666-7568(23)00037-5

130. Nicholson T, Dhaliwal A, Quinlan JI, Allen SL, Williams FR, Hazeldine J, McGee KC, Sullivan J, Breen L, Elsharkawy AM, Armstrong MJ, Jones SW, Greig CA, Lord JM (2024) Accelerated aging of skeletal muscle and the immune system in patients with chronic liver disease. Exp Mol Med 56:1667–1681. 10.1038/s12276-024-01287-y

131. Li Z, Zong X, Li D, He Y, Tang J, Hu M, Chen X (2023) Epigenetic clock analysis of blood samples in drug-naive first-episode schizophrenia patients. BMC Psychiatry 23:45. 10.1186/s12888-023-04533-1

132. Bejaoui Y, Humaira Amanullah F, Saad M, Taleb S, Bradic M, Megarbane A, Ait Hssain A, Abi Khalil C, El Hajj N (2023) Epigenetic age acceleration in surviving versus deceased COVID-19 patients with acute respiratory distress syndrome following hospitalization. Clinical Epigenetics 15:186. 10.1186/s13148-023-01597-4

133. Higgins-Chen AT, Boks MP, Vinkers CH, Kahn RS, Levine ME (2020) Schizophrenia and Epigenetic Aging Biomarkers: Increased Mortality, Reduced Cancer Risk, and Unique Clozapine Effects. Biol Psychiatry 88:224–235. 10.1016/j.biopsych.2020.01.025

134. Shinko Y, Okazaki S, Otsuka I, Horai T, Kim S, Tanifuji T, Hishimoto A (2022) Accelerated epigenetic age and shortened telomere length based on DNA methylation in Nicolaides–Baraitser syndrome. Mol Genet Genomic Med 10:e1876. 10.1002/mgg3.1876

135. McCarthy K, O’Halloran AM, Fallon P, Kenny RA, McCrory C (2023) Metabolic syndrome accelerates epigenetic ageing in older adults: Findings from The Irish Longitudinal Study on Ageing (TILDA). Experimental Gerontology 183:112314. 10.1016/j.exger.2023.112314

136. Protsenko E, Yang R, Nier B, Reus V, Hammamieh R, Rampersaud R, Wu GWY, Hough CM, Epel E, Prather AA, Jett M, Gautam A, Mellon SH, Wolkowitz OM (2021) “GrimAge,” an epigenetic predictor of mortality, is accelerated in major depressive disorder. Transl Psychiatry 11:1–9. 10.1038/s41398-021-01302-0

137. Lima CNC, Suchting R, Scaini G, Cuellar VA, Favero-Campbell AD, Walss-Bass C, Soares JC, Quevedo J, Fries GR (2022) Epigenetic GrimAge acceleration and cognitive impairment in bipolar disorder. Eur Neuropsychopharmacol 62:10–21. 10.1016/j.euroneuro.2022.06.007

138. García-delaTorre P, Rivero-Segura NA, Sánchez-García S, Becerril-Rojas K, Sandoval-Rodriguez FE, Castro-Morales D, Cruz-Lopez M, Vazquez-Moreno M, Rincón-Heredia R, Ramirez-Garcia P, Gomez-Verjan JC (2024) GrimAge is elevated in older adults with mild COVID-19 an exploratory analysis. GeroScience 46:3511–3524. 10.1007/s11357-024-01095-2

139. Wikström Shemer D, Mostafaei S, Tang B, Pedersen NL, Karlsson IK, Fall T, Hägg S (2024) Associations between epigenetic aging and diabetes mellitus in a Swedish longitudinal study. GeroScience 46:5003–5014. 10.1007/s11357-024-01252-7

140. Miao K, Hong X, Cao W, Lv J, Yu C, Huang T, Sun D, Liao C, Pang Y, Hu R, Pang Z, Yu M, Wang H, Wu X, Liu Y, Gao W, Li L (2024) Association between epigenetic age and type 2 diabetes mellitus or glycemic traits: A longitudinal twin study. Aging Cell 23:e14175. 10.1111/acel.14175

141. Föhr T, Hendrix A, Kankaanpää A, Laakkonen EK, Kujala U, Pietiläinen KH, Lehtimäki T, Kähönen M, Raitakari O, Wang X, Kaprio J, Ollikainen M, Sillanpää E (2024) Metabolic syndrome and epigenetic aging: a twin study. Int J Obes 48:778–787. 10.1038/s41366-024-01466-x

142. Aroke EN, Wiggins AM, Hobson JM, Srinivasasainagendra V, Quinn TL, Kottae P, Tiwari HK, Sorge RE, Goodin BR (2023) The pace of biological aging helps explain the association between insomnia and chronic low back pain. Mol Pain 19:17448069231210648. 10.1177/17448069231210648

143. Thomas A, Ryan CP, Caspi A, Liu Z, Moffitt TE, Sugden K, Zhou J, Belsky DW, Gu Y (2024) Diet, Pace of Biological Aging, and Risk of Dementia in the Framingham Heart Study. Ann Neurol 95:1069–1079. 10.1002/ana.26900

144. Whitman ET, Ryan CP, Abraham WC, Addae A, Corcoran DL, Elliott ML, Hogan S, Ireland D, Keenan R, Knodt AR, Melzer TR, Poulton R, Ramrakha S, Sugden K, Williams BS, Zhou J, Hariri AR, Belsky DW, Moffitt TE, Caspi A (2024) A blood biomarker of the pace of aging is associated with brain structure: replication across three cohorts. Neurobiology of Aging 136:23–33. 10.1016/j.neurobiolaging.2024.01.008

145. Sugden K, Caspi A, Elliott ML, Bourassa KJ, Chamarti K, Corcoran DL, Hariri AR, Houts RM, Kothari M, Kritchevsky S, Kuchel GA, Mill JS, Williams BS, Belsky DW, Moffitt TE, for the Alzheimer’s Disease Neuroimaging Initiative* (2022) Association of Pace of Aging Measured by Blood-Based DNA Methylation With Age-Related Cognitive Impairment and Dementia. Neurology 99:e1402–e1413. 10.1212/WNL.0000000000200898

