## Supplementary Figures for "EpImAge: An Epigenetic-Immune Clock for Disease-Associated Biological Aging"

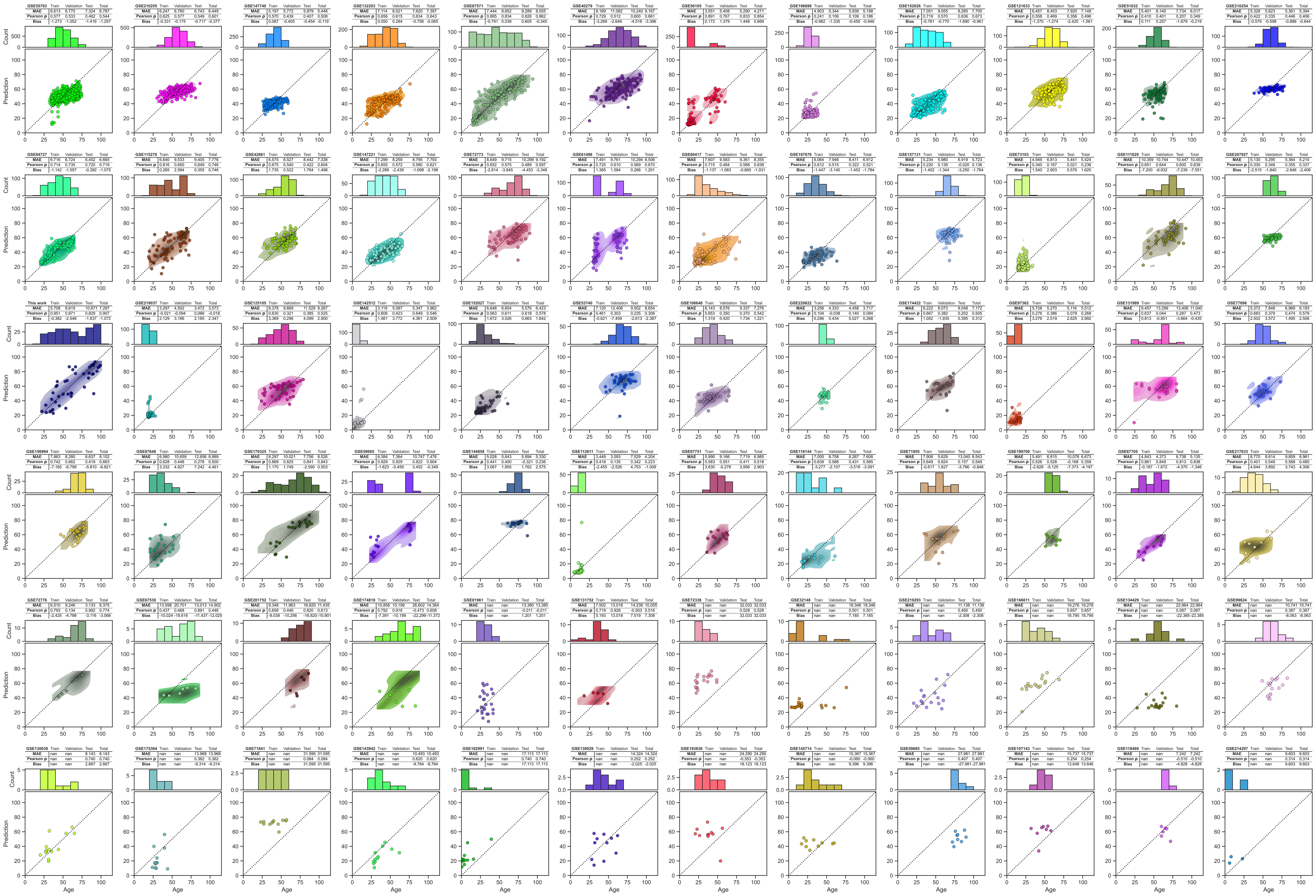


Supplementary Figure S1. EpImAge results for all considered datasets. For each dataset, the relationship between real and predicted age is presented, KDE corresponds to train-validation data, scatter corresponds to test data (if the dataset is too small, it only participates in the test data). The table for each dataset shows MAE values and Pearson correlation coefficient total and separately for train, validation, test data.

| | | | | | Chapter 1 | | Chapter 2 | | | | Chapter 4 | | Chapter 5 | | | | | Chapter 6 | | | | | | Chapter 8 | | | | | | | 11 | 12 | Chapter 13 | | Chapter 15 | | Chapter 16 | Chapter 20 | | Chapter 25 | |
|---|---|---|---|---|---|---|---|---|---|---|---|---|---|---|---|---|---|---|---|---|---|---|---|---|---|---|---|---|---|---|---|---|---|---|---|---|---|---|---|---|---|
| Clocks | Year | Total Pearson ρ | Total MAE | Passed ICD-11 | 1B10 | 1C60-1C62 | 2B90 | 2B91 | 2B92 | 2C60-2C6Z | 4A40.0 | 4A44.2 | 5A02.0 | 5A10 | 5A40 | 5A61.0 | 5B81 | 6A20 | 6A21 | 6A23 | 6A70-6A71 | 6D71 | 6D83 | 8A00.0 | 8A00.10 | 8A20 | 8A40 | 8A40.0 | 8A40.2 | 8E00 | BA52, BD40 | CB00 | DD70 | DD71 | FA20 | FB83.1 | GB61.5 | LD2B | LD2F.1Y | RA01 | RA02 |
| This work | | 0.850 | 7.038 | 25/76 | 0/3 | 5/8 | 0/1 | 0/1 | 0/1 | 0/1 | 1/1 | 0/2 | 0/1 | 1/1 | 0/1 | 2/2 | 0/3 | 2/3 | 0/1 | 2/2 | 1/1 | 0/1 | 0/1 | 0/2 | 0/1 | 0/3 | 0/2 | 0/2 | 0/2 | 1/1 | 0/3 | 1/1 | 1/4 | 1/2 | 1/5 | 0/1 | 1/1 | 0/2 | 2/2 | 2/6 | 1/1 |
| Hannum | 2013 | 0.896 | 6.820 | 14/76 | 0/3 | 3/8 | 1/1 | 0/1 | 0/1 | 0/1 | 0/1 | 0/2 | 0/1 | 0/1 | 0/1 | 0/2 | 0/3 | 1/3 | 0/1 | 2/2 | 1/1 | 0/1 | 0/1 | 0/2 | 1/1 | 0/3 | 0/2 | 0/2 | 0/2 | 0/1 | 0/3 | 1/1 | 2/4 | 1/2 | 0/5 | 0/1 | 1/1 | 0/2 | 0/2 | 0/6 | 0/1 |
| Horvath | 2013 | 0.931 | 5.425 | 9/76 | 0/3 | 5/8 | 1/1 | 0/1 | 0/1 | 0/1 | 0/1 | 0/2 | 0/1 | 0/1 | 0/1 | 0/2 | 0/3 | 1/3 | 0/1 | 1/2 | 0/1 | 0/1 | 0/1 | 0/2 | 0/1 | 0/3 | 0/2 | 0/2 | 0/2 | 0/1 | 0/3 | 0/1 | 0/4 | 0/2 | 0/5 | 0/1 | 1/1 | 0/2 | 0/2 | 0/6 | 0/1 |
| Lin | 2016 | 0.899 | 6.842 | 10/76 | 0/3 | 6/8 | 0/1 | 0/1 | 0/1 | 0/1 | 0/1 | 0/2 | 0/1 | 0/1 | 0/1 | 0/2 | 0/3 | 1/3 | 0/1 | 1/2 | 0/1 | 0/1 | 0/1 | 0/2 | 1/1 | 0/3 | 0/2 | 0/2 | 0/2 | 0/1 | 0/3 | 0/1 | 0/4 | 0/2 | 0/5 | 0/1 | 0/1 | 0/2 | 0/2 | 0/6 | 1/1 |
| epiTOC1 | 2016 | | | 9/76 | 0/3 | 5/8 | 0/1 | 0/1 | 0/1 | 0/1 | 0/1 | 0/2 | 0/1 | 0/1 | 0/1 | 0/2 | 1/3 | 1/3 | 0/1 | 1/2 | 0/1 | 0/1 | 0/1 | 0/2 | 0/1 | 0/3 | 0/2 | 0/2 | 0/2 | 0/1 | 0/3 | 0/1 | 0/4 | 0/2 | 0/5 | 0/1 | 0/1 | 0/2 | 0/2 | 0/6 | 1/1 |
| ZhangMortality | 2017 | | | 24/76 | 0/3 | 3/8 | 1/1 | 0/1 | 0/1 | 0/1 | 0/1 | 1/2 | 0/1 | 0/1 | 0/1 | 0/2 | 0/3 | 3/3 | 1/1 | 0/2 | 0/1 | 0/1 | 1/1 | 2/2 | 0/1 | 1/3 | 0/2 | 0/2 | 0/2 | 1/1 | 1/3 | 1/1 | 3/4 | 1/2 | 3/5 | 0/1 | 0/1 | 0/2 | 0/2 | 1/6 | 0/1 |
| DNAmPhenoAge | 2018 | 0.912 | 8.107 | 23/76 | 1/3 | 5/8 | 0/1 | 0/1 | 0/1 | 0/1 | 0/1 | 0/2 | 0/1 | 0/1 | 0/1 | 0/2 | 0/3 | 2/3 | 1/1 | 2/2 | 1/1 | 0/1 | 0/1 | 0/2 | 0/1 | 0/3 | 0/2 | 0/2 | 0/2 | 1/1 | 1/3 | 1/1 | 2/4 | 1/2 | 2/5 | 0/1 | 1/1 | 1/2 | 0/2 | 1/6 | 0/1 |
| SkinAndBlood | 2018 | 0.961 | 3.809 | 16/76 | 0/3 | 5/8 | 0/1 | 0/1 | 0/1 | 0/1 | 1/1 | 0/2 | 0/1 | 0/1 | 0/1 | 1/2 | 0/3 | 1/3 | 1/1 | 2/2 | 0/1 | 0/1 | 0/1 | 0/2 | 0/1 | 0/3 | 0/2 | 0/2 | 0/2 | 0/1 | 0/3 | 0/1 | 0/4 | 0/2 | 0/5 | 0/1 | 1/1 | 1/2 | 0/2 | 2/6 | 1/1 |
| GrimAge | 2019 | 0.467 | 16.869 | 22/76 | 0/3 | 3/8 | 0/1 | 0/1 | 0/1 | 0/1 | 1/1 | 0/2 | 0/1 | 0/1 | 0/1 | 2/2 | 0/3 | 2/3 | 0/1 | 2/2 | 1/1 | 0/1 | 0/1 | 0/2 | 0/1 | 0/3 | 1/2 | 0/2 | 0/2 | 1/1 | 1/3 | 1/1 | 2/4 | 0/2 | 1/5 | 0/1 | 1/1 | 0/2 | 1/2 | 1/6 | 1/1 |
| DNAmTL | 2019 | | | 17/76 | 0/3 | 5/8 | 1/1 | 0/1 | 0/1 | 0/1 | 0/1 | 0/2 | 0/1 | 0/1 | 0/1 | 0/2 | 1/3 | 1/3 | 0/1 | 0/2 | 0/1 | 0/1 | 0/1 | 0/2 | 0/1 | 0/3 | 0/2 | 0/2 | 0/2 | 0/1 | 3/3 | 1/1 | 2/4 | 0/2 | 1/5 | 0/1 | 0/1 | 1/2 | 0/2 | 1/6 | 0/1 |
| ZhangEN | 2019 | 0.976 | 6.805 | 6/76 | 0/3 | 3/8 | 0/1 | 0/1 | 0/1 | 0/1 | 0/1 | 0/2 | 0/1 | 0/1 | 0/1 | 0/2 | 0/3 | 1/3 | 1/1 | 0/2 | 0/1 | 0/1 | 0/1 | 0/2 | 0/1 | 0/3 | 0/2 | 0/2 | 0/2 | 0/1 | 0/3 | 0/1 | 1/4 | 0/2 | 0/5 | 0/1 | 0/1 | 0/2 | 0/2 | 0/6 | 0/1 |
| ZhangBLUP | 2019 | 0.979 | 2.606 | 9/76 | 0/3 | 4/8 | 0/1 | 0/1 | 0/1 | 0/1 | 0/1 | 0/2 | 0/1 | 0/1 | 0/1 | 0/2 | 0/3 | 1/3 | 0/1 | 0/2 | 1/1 | 0/1 | 0/1 | 0/2 | 0/1 | 0/3 | 0/2 | 0/2 | 0/2 | 0/1 | 0/3 | 0/1 | 1/4 | 0/2 | 0/5 | 0/1 | 1/1 | 1/2 | 0/2 | 0/6 | 0/1 |
| Han | 2020 | 0.925 | 5.226 | 13/76 | 0/3 | 6/8 | 1/1 | 0/1 | 0/1 | 0/1 | 0/1 | 0/2 | 0/1 | 0/1 | 0/1 | 1/2 | 1/3 | 1/3 | 0/1 | 1/2 | 0/1 | 0/1 | 0/1 | 0/2 | 0/1 | 0/3 | 0/2 | 0/2 | 0/2 | 0/1 | 1/3 | 0/1 | 0/4 | 0/2 | 0/5 | 0/1 | 0/1 | 1/2 | 0/2 | 0/6 | 0/1 |
| DunedinPACE | 2022 | | | 36/76 | 1/3 | 5/8 | 1/1 | 0/1 | 0/1 | 0/1 | 0/1 | 1/2 | 1/1 | 0/1 | 0/1 | 2/2 | 2/3 | 3/3 | 1/1 | 1/2 | 0/1 | 0/1 | 1/1 | 0/2 | 0/1 | 0/3 | 1/2 | 0/2 | 0/2 | 1/1 | 1/3 | 1/1 | 4/4 | 1/2 | 4/5 | 0/1 | 1/1 | 1/2 | 0/2 | 2/6 | 0/1 |
| AltumAge | 2022 | 0.920 | 5.439 | 12/76 | 0/3 | 4/8 | 0/1 | 0/1 | 0/1 | 0/1 | 0/1 | 0/2 | 0/1 | 0/1 | 0/1 | 0/2 | 0/3 | 0/3 | 0/1 | 2/2 | 0/1 | 0/1 | 0/1 | 0/2 | 0/1 | 0/3 | 0/2 | 0/2 | 0/2 | 1/1 | 2/3 | 0/1 | 0/4 | 0/2 | 0/5 | 0/1 | 1/1 | 0/2 | 0/2 | 1/6 | 1/1 |
| PCHannum | 2022 | 0.914 | 8.238 | 19/76 | 0/3 | 6/8 | 1/1 | 0/1 | 0/1 | 0/1 | 0/1 | 0/2 | 0/1 | 0/1 | 0/1 | 0/2 | 0/3 | 2/3 | 0/1 | 1/2 | 1/1 | 0/1 | 0/1 | 0/2 | 1/1 | 0/3 | 0/2 | 0/2 | 0/2 | 1/1 | 0/3 | 1/1 | 2/4 | 1/2 | 1/5 | 0/1 | 1/1 | 0/2 | 0/2 | 0/6 | 0/1 |
| PCHorvath | 2022 | 0.904 | 6.827 | 11/76 | 0/3 | 6/8 | 0/1 | 0/1 | 0/1 | 0/1 | 0/1 | 0/2 | 0/1 | 0/1 | 0/1 | 0/2 | 0/3 | 1/3 | 0/1 | 1/2 | 1/1 | 0/1 | 0/1 | 0/2 | 0/1 | 0/3 | 0/2 | 0/2 | 0/2 | 0/1 | 0/3 | 0/1 | 0/4 | 0/2 | 0/5 | 0/1 | 1/1 | 0/2 | 0/2 | 0/6 | 1/1 |
| PCPhenoAge | 2022 | 0.895 | 6.478 | 31/76 | 2/3 | 6/8 | 0/1 | 0/1 | 0/1 | 0/1 | 0/1 | 0/2 | 0/1 | 0/1 | 0/1 | 0/2 | 0/3 | 2/3 | 1/1 | 2/2 | 1/1 | 0/1 | 1/1 | 2/2 | 1/1 | 0/3 | 2/2 | 0/2 | 0/2 | 1/1 | 0/3 | 1/1 | 2/4 | 1/2 | 1/5 | 0/1 | 1/1 | 1/2 | 0/2 | 2/6 | 1/1 |
| HRSInCHPhenoAge | 2022 | 0.913 | 6.621 | 34/76 | 2/3 | 6/8 | 1/1 | 0/1 | 0/1 | 0/1 | 0/1 | 0/2 | 0/1 | 0/1 | 0/1 | 0/2 | 1/3 | 3/3 | 1/1 | 2/2 | 1/1 | 0/1 | 0/1 | 2/2 | 1/1 | 0/3 | 0/2 | 0/2 | 0/2 | 1/1 | 1/3 | 1/1 | 2/4 | 1/2 | 3/5 | 0/1 | 1/1 | 1/2 | 0/2 | 2/6 | 1/1 |
| PCSkinAndBlood | 2022 | 0.897 | 6.627 | 11/76 | 0/3 | 6/8 | 0/1 | 0/1 | 0/1 | 0/1 | 0/1 | 0/2 | 0/1 | 0/1 | 0/1 | 0/2 | 0/3 | 1/3 | 0/1 | 1/2 | 1/1 | 0/1 | 0/1 | 0/2 | 0/1 | 0/3 | 0/2 | 0/2 | 0/2 | 1/1 | 0/3 | 0/1 | 0/4 | 0/2 | 0/5 | 0/1 | 1/1 | 0/2 | 0/2 | 0/6 | 0/1 |
| PCGrimAge | 2022 | 0.558 | 24.101 | 20/76 | 0/3 | 3/8 | 0/1 | 0/1 | 0/1 | 0/1 | 1/1 | 0/2 | 0/1 | 0/1 | 0/1 | 2/2 | 0/3 | 2/3 | 0/1 | 2/2 | 1/1 | 0/1 | 0/1 | 0/2 | 0/1 | 0/3 | 0/2 | 0/2 | 0/2 | 1/1 | 0/3 | 1/1 | 2/4 | 1/2 | 1/5 | 0/1 | 1/1 | 0/2 | 1/2 | 1/6 | 1/1 |
| PCDNAmTL | 2022 | | | 19/76 | 0/3 | 5/8 | 1/1 | 0/1 | 0/1 | 0/1 | 0/1 | 0/2 | 0/1 | 0/1 | 0/1 | 0/2 | 1/3 | 1/3 | 0/1 | 0/2 | 0/1 | 0/1 | 0/1 | 1/2 | 1/1 | 0/3 | 0/2 | 0/2 | 0/2 | 0/1 | 3/3 | 0/1 | 2/4 | 1/2 | 1/5 | 0/1 | 0/1 | 1/2 | 0/2 | 1/6 | 0/1 |
| GrimAge2 | 2022 | 0.529 | 19.419 | 27/76 | 0/3 | 3/8 | 0/1 | 0/1 | 0/1 | 0/1 | 1/1 | 0/2 | 0/1 | 0/1 | 0/1 | 2/2 | 0/3 | 2/3 | 0/1 | 2/2 | 1/1 | 0/1 | 0/1 | 0/2 | 0/1 | 0/3 | 1/2 | 0/2 | 0/2 | 1/1 | 1/3 | 1/1 | 3/4 | 1/2 | 1/5 | 0/1 | 1/1 | 0/2 | 2/2 | 3/6 | 1/1 |
| DNAmFitAge | 2023 | 0.888 | 14.558 | 12/76 | 0/3 | 6/8 | 0/1 | 0/1 | 0/1 | 0/1 | 1/1 | 0/2 | 0/1 | 0/1 | 0/1 | 0/2 | 0/3 | 1/3 | 0/1 | 2/2 | 1/1 | 0/1 | 0/1 | 0/2 | 0/1 | 0/3 | 0/2 | 0/2 | 0/2 | 1/1 | 0/3 | 0/1 | 0/4 | 0/2 | 0/5 | 0/1 | 0/1 | 0/2 | 0/2 | 0/6 | 0/1 |
| ENCen40 | 2023 | 0.959 | 5.345 | 14/76 | 0/3 | 6/8 | 0/1 | 0/1 | 0/1 | 0/1 | 0/1 | 0/2 | 0/1 | 0/1 | 0/1 | 0/2 | 0/3 | 1/3 | 0/1 | 2/2 | 0/1 | 0/1 | 0/1 | 0/2 | 1/1 | 0/3 | 0/2 | 0/2 | 0/2 | 0/1 | 0/3 | 0/1 | 1/4 | 0/2 | 0/5 | 0/1 | 0/1 | 1/2 | 0/2 | 1/6 | 1/1 |
| YingAdaptAge | 2024 | 0.559 | 17.244 | 8/76 | 0/3 | 3/8 | 0/1 | 0/1 | 0/1 | 1/1 | 0/1 | 0/2 | 0/1 | 0/1 | 0/1 | 0/2 | 0/3 | 1/3 | 0/1 | 2/2 | 0/1 | 0/1 | 0/1 | 0/2 | 0/1 | 0/3 | 0/2 | 0/2 | 0/2 | 0/1 | 0/3 | 0/1 | 1/4 | 0/2 | 0/5 | 0/1 | 0/1 | 0/2 | 0/2 | 0/6 | 0/1 |
| YingCausAge | 2024 | 0.589 | 23.329 | 13/76 | 0/3 | 6/8 | 0/1 | 0/1 | 0/1 | 0/1 | 0/1 | 0/2 | 0/1 | 0/1 | 0/1 | 0/2 | 0/3 | 2/3 | 0/1 | 1/2 | 0/1 | 0/1 | 0/1 | 0/2 | 1/1 | 0/3 | 0/2 | 0/2 | 0/2 | 0/1 | 0/3 | 0/1 | 1/4 | 0/2 | 0/5 | 0/1 | 0/1 | 1/2 | 0/2 | 0/6 | 1/1 |
| YingDamAge | 2024 | 0.467 | 18.330 | 11/76 | 0/3 | 1/8 | 0/1 | 0/1 | 0/1 | 0/1 | 0/1 | 0/2 | 0/1 | 0/1 | 0/1 | 0/2 | 0/3 | 1/3 | 0/1 | 2/2 | 0/1 | 0/1 | 0/1 | 0/2 | 0/1 | 0/3 | 0/2 | 0/2 | 0/2 | 0/1 | 0/3 | 1/1 | 1/4 | 0/2 | 0/5 | 0/1 | 1/1 | 0/2 | 0/2 | 3/6 | 1/1 |
| StocH | 2024 | 0.702 | 10.455 | 6/76 | 0/3 | 4/8 | 0/1 | 0/1 | 0/1 | 0/1 | 0/1 | 0/2 | 0/1 | 0/1 | 0/1 | 0/2 | 0/3 | 0/3 | 0/1 | 2/2 | 0/1 | 0/1 | 0/1 | 0/2 | 0/1 | 0/3 | 0/2 | 0/2 | 0/2 | 0/1 | 0/3 | 0/1 | 0/4 | 0/2 | 0/5 | 0/1 | 0/1 | 0/2 | 0/2 | 0/6 | 0/1 |
| StocP | 2024 | 0.755 | 10.122 | 7/76 | 0/3 | 4/8 | 0/1 | 0/1 | 0/1 | 0/1 | 0/1 | 0/2 | 0/1 | 0/1 | 0/1 | 0/2 | 0/3 | 0/3 | 0/1 | 1/2 | 0/1 | 0/1 | 0/1 | 0/2 | 0/1 | 0/3 | 0/2 | 0/2 | 0/2 | 0/1 | 0/3 | 0/1 | 0/4 | 0/2 | 0/5 | 0/1 | 1/1 | 0/2 | 1/2 | 0/6 | 0/1 |
| StocZ | 2024 | 0.942 | 11.687 | 7/76 | 0/3 | 5/8 | 0/1 | 0/1 | 0/1 | 0/1 | 0/1 | 0/2 | 0/1 | 0/1 | 0/1 | 0/2 | 1/3 | 0/3 | 0/1 | 0/2 | 0/1 | 0/1 | 0/1 | 1/2 | 0/1 | 0/3 | 0/2 | 0/2 | 0/2 | 0/1 | 0/3 | 0/1 | 0/4 | 0/2 | 0/5 | 0/1 | 0/1 | 0/2 | 0/2 | 0/6 | 0/1 |
| stemTOC | 2024 | | | 6/76 | 0/3 | 5/8 | 0/1 | 0/1 | 0/1 | 0/1 | 0/1 | 0/2 | 0/1 | 0/1 | 0/1 | 0/2 | 0/3 | 0/3 | 0/1 | 1/2 | 0/1 | 0/1 | 0/1 | 0/2 | 0/1 | 0/3 | 0/2 | 0/2 | 0/2 | 0/1 | 0/3 | 0/1 | 0/4 | 0/2 | 0/5 | 0/1 | 0/1 | 0/2 | 0/2 | 0/6 | 0/1 |
| RetroelementAgeV1 | 2024 | 0.780 | 27.900 | 11/76 | 0/3 | 6/8 | 0/1 | 0/1 | 0/1 | 0/1 | 1/1 | 0/2 | 0/1 | 0/1 | 0/1 | 0/2 | 0/3 | 0/3 | 0/1 | 2/2 | 1/1 | 0/1 | 0/1 | 0/2 | 0/1 | 0/3 | 0/2 | 0/2 | 0/2 | 0/1 | 0/3 | 0/1 | 0/4 | 0/2 | 0/5 | 0/1 | 0/1 | 1/2 | 0/2 | 0/6 | 0/1 |
| RetroelementAgeV2 | 2024 | 0.806 | 12.017 | 9/76 | 0/3 | 2/8 | 0/1 | 0/1 | 0/1 | 0/1 | 1/1 | 0/2 | 0/1 | 0/1 | 0/1 | 1/2 | 0/3 | 0/3 | 0/1 | 2/2 | 1/1 | 0/1 | 0/1 | 0/2 | 0/1 | 0/3 | 0/2 | 0/2 | 0/2 | 0/1 | 0/3 | 0/1 | 0/4 | 0/2 | 0/5 | 0/1 | 0/1 | 1/2 | 1/2 | 0/6 | 0/1 |
| IntrinClock | 2024 | 0.960 | 3.719 | 7/76 | 0/3 | 2/8 | 0/1 | 0/1 | 0/1 | 0/1 | 0/1 | 0/2 | 0/1 | 0/1 | 0/1 | 0/2 | 0/3 | 1/3 | 0/1 | 0/2 | 0/1 | 0/1 | 0/1 | 0/2 | 0/1 | 0/3 | 0/2 | 0/2 | 0/2 | 0/1 | 0/3 | 0/1 | 0/4 | 0/2 | 0/5 | 0/1 | 1/1 | 1/2 | 0/2 | 1/6 | 1/1 |

Suppllementary Figure S2. Detailed performance results of different epigenetic clocks (including the ones presented in this paper). The first column gives the names of the clocks. For each epigenetic clock model, the year of development (clocks are ordered by the chronology of their release), common metrics - Pearson correlation coefficient and MAE, and the number of disease sensitivity tests passed (Mann-Whitney p-value < 0.05) are given both overall and for each ICD-11 disease code.

| | | Chapter 1 | | Chapter 2 | | | | Chapter 4 | | Chapter 5 | | | | | Chapter 6 | | | | |
|---|---|---|---|---|---|---|---|---|---|---|---|---|---|---|---|---|---|---|---|
| Immunomarker | Passed ICD-11 | 1B10 | 1C60-1C62 | 2B90 | 2B91 | 2B92 | 2C60-2C6Z | 4A40.0 | 4A44.2 | 5A02.0 | 5A10 | 5A40 | 5A61.0 | 5B81 | 6A20 | 6A21 | 6A23 | 6A70-6A71 | 6D71 |
| CXCL9 | 25/76 | 0/3 | 2/8 | 0/1 | 0/1 | 0/1 | 1/1 | 0/1 | 0/2 | 1/1 | 0/1 | 0/1 | 0/2 | 2/3 | 3/3 | 1/1 | 1/2 | 0/1 | 0/1 |
| CCL11 | 2/76 | 0/3 | 0/8 | 0/1 | 0/1 | 0/1 | 0/1 | 0/1 | 0/2 | 0/1 | 0/1 | 1/1 | 0/2 | 0/3 | 0/3 | 0/1 | 0/2 | 0/1 | 0/1 |
| IL27 | 14/76 | 0/3 | 2/8 | 0/1 | 0/1 | 0/1 | 1/1 | 0/1 | 0/2 | 0/1 | 0/1 | 0/1 | 0/2 | 2/3 | 1/3 | 0/1 | 0/2 | 0/1 | 0/1 |
| IL5 | 2/76 | 0/3 | 0/8 | 0/1 | 0/1 | 0/1 | 0/1 | 0/1 | 0/2 | 0/1 | 0/1 | 0/1 | 0/2 | 0/3 | 0/3 | 0/1 | 0/2 | 0/1 | 0/1 |
| CSF1 | 11/76 | 0/3 | 1/8 | 0/1 | 0/1 | 0/1 | 0/1 | 1/1 | 0/2 | 1/1 | 0/1 | 0/1 | 0/2 | 0/3 | 1/3 | 0/1 | 1/2 | 0/1 | 0/1 |
| CCL2 | 15/76 | 1/3 | 2/8 | 0/1 | 0/1 | 0/1 | 0/1 | 0/1 | 0/2 | 0/1 | 0/1 | 0/1 | 0/2 | 2/3 | 3/3 | 0/1 | 0/2 | 0/1 | 0/1 |
| IL1B | 11/76 | 0/3 | 4/8 | 0/1 | 0/1 | 0/1 | 1/1 | 0/1 | 0/2 | 0/1 | 0/1 | 0/1 | 0/2 | 1/3 | 1/3 | 0/1 | 1/2 | 0/1 | 0/1 |
| IL6 | 24/76 | 0/3 | 3/8 | 1/1 | 0/1 | 0/1 | 0/1 | 0/1 | 0/2 | 0/1 | 0/1 | 0/1 | 2/2 | 1/3 | 3/3 | 1/1 | 1/2 | 0/1 | 0/1 |
| GCSF | 6/76 | 0/3 | 1/8 | 0/1 | 0/1 | 0/1 | 1/1 | 0/1 | 0/2 | 0/1 | 0/1 | 0/1 | 0/2 | 0/3 | 0/3 | 0/1 | 0/2 | 0/1 | 0/1 |
| CXCL10 | 17/76 | 0/3 | 4/8 | 0/1 | 0/1 | 0/1 | 1/1 | 0/1 | 0/2 | 0/1 | 0/1 | 1/1 | 1/2 | 0/3 | 3/3 | 0/1 | 0/2 | 0/1 | 0/1 |
| VEGFA | 7/76 | 0/3 | 3/8 | 0/1 | 0/1 | 0/1 | 0/1 | 0/1 | 0/2 | 0/1 | 0/1 | 0/1 | 0/2 | 0/3 | 0/3 | 0/1 | 0/2 | 0/1 | 0/1 |
| TNF | 8/76 | 0/3 | 2/8 | 0/1 | 0/1 | 0/1 | 0/1 | 0/1 | 0/2 | 0/1 | 0/1 | 0/1 | 0/2 | 0/3 | 0/3 | 0/1 | 0/2 | 0/1 | 0/1 |
| PDGFB | 4/76 | 0/3 | 1/8 | 1/1 | 0/1 | 0/1 | 0/1 | 0/1 | 0/2 | 0/1 | 0/1 | 0/1 | 0/2 | 0/3 | 0/3 | 0/1 | 0/2 | 0/1 | 0/1 |
| IL8 | 9/76 | 0/3 | 2/8 | 0/1 | 0/1 | 0/1 | 1/1 | 0/1 | 0/2 | 0/1 | 0/1 | 0/1 | 1/2 | 0/3 | 1/3 | 0/1 | 0/2 | 0/1 | 0/1 |
| PDGFA | 0/76 | 0/3 | 0/8 | 0/1 | 0/1 | 0/1 | 0/1 | 0/1 | 0/2 | 0/1 | 0/1 | 0/1 | 0/2 | 0/3 | 0/3 | 0/1 | 0/2 | 0/1 | 0/1 |
| IL12Bp40 | 5/76 | 0/3 | 1/8 | 0/1 | 0/1 | 0/1 | 0/1 | 1/1 | 0/2 | 0/1 | 0/1 | 0/1 | 0/2 | 0/3 | 0/3 | 0/1 | 1/2 | 0/1 | 0/1 |
| IL15 | 17/76 | 0/3 | 5/8 | 0/1 | 0/1 | 0/1 | 0/1 | 0/1 | 1/2 | 1/1 | 0/1 | 0/1 | 1/2 | 0/3 | 1/3 | 0/1 | 0/2 | 1/1 | 0/1 |
| CXCL1 | 7/76 | 0/3 | 1/8 | 0/1 | 0/1 | 0/1 | 1/1 | 0/1 | 0/2 | 0/1 | 0/1 | 0/1 | 0/2 | 0/3 | 0/3 | 0/1 | 1/2 | 0/1 | 0/1 |
| CCL4 | 3/76 | 0/3 | 1/8 | 0/1 | 0/1 | 0/1 | 0/1 | 0/1 | 0/2 | 0/1 | 0/1 | 0/1 | 0/2 | 0/3 | 0/3 | 0/1 | 0/2 | 0/1 | 0/1 |
| IFNA2 | 5/76 | 0/3 | 0/8 | 0/1 | 0/1 | 0/1 | 1/1 | 0/1 | 0/2 | 0/1 | 0/1 | 0/1 | 0/2 | 2/3 | 1/3 | 0/1 | 0/2 | 0/1 | 0/1 |
| IL13 | 7/76 | 0/3 | 0/8 | 1/1 | 0/1 | 0/1 | 1/1 | 0/1 | 0/2 | 0/1 | 0/1 | 0/1 | 1/2 | 0/3 | 0/3 | 0/1 | 0/2 | 0/1 | 0/1 |
| FLT3L | 11/76 | 0/3 | 5/8 | 0/1 | 0/1 | 0/1 | 0/1 | 0/1 | 0/2 | 0/1 | 0/1 | 0/1 | 0/2 | 0/3 | 0/3 | 0/1 | 1/2 | 0/1 | 0/1 |
| CD40LG | 4/76 | 0/3 | 0/8 | 0/1 | 0/1 | 0/1 | 0/1 | 1/1 | 0/2 | 0/1 | 0/1 | 0/1 | 0/2 | 0/3 | 1/3 | 0/1 | 0/2 | 0/1 | 0/1 |
| CCL22 | 11/76 | 0/3 | 0/8 | 0/1 | 0/1 | 0/1 | 1/1 | 0/1 | 1/2 | 0/1 | 0/1 | 0/1 | 1/2 | 0/3 | 1/3 | 0/1 | 0/2 | 0/1 | 0/1 |

| | Chapter 6 | Chapter 8 | | | | | | | 11 | 12 | Chapter 13 | | Chapter 15 | | Chapter 16 | Chapter 20 | | Chapter 25 | |
|---|---|---|---|---|---|---|---|---|---|---|---|---|---|---|---|---|---|---|---|
| Immunomarker | 6D83 | 8A00.0 | 8A00.10 | 8A20 | 8A40 | 8A40.0 | 8A40.2 | 8E00 | BA52, BD40 | CB00 | DD70 | DD71 | FA20 | FB83.1 | GB61.5 | LD2B | LD2F.1Y | RA01 | RA02 |
| CXCL9 | 0/1 | 1/2 | 0/1 | 0/3 | 1/2 | 0/2 | 0/2 | 0/1 | 1/3 | 1/1 | 2/4 | 1/2 | 1/5 | 0/1 | 1/1 | 1/2 | 0/2 | 3/6 | 1/1 |
| CCL11 | 0/1 | 0/2 | 0/1 | 0/3 | 0/2 | 0/2 | 0/2 | 0/1 | 0/3 | 0/1 | 0/4 | 0/2 | 0/5 | 0/1 | 1/1 | 0/2 | 0/2 | 0/6 | 0/1 |
| IL27 | 0/1 | 2/2 | 0/1 | 0/3 | 0/2 | 0/2 | 0/2 | 0/1 | 0/3 | 0/1 | 1/4 | 1/2 | 1/5 | 0/1 | 1/1 | 0/2 | 1/2 | 1/6 | 0/1 |
| IL5 | 0/1 | 0/2 | 0/1 | 0/3 | 0/2 | 0/2 | 0/2 | 0/1 | 0/3 | 0/1 | 0/4 | 0/2 | 1/5 | 0/1 | 0/1 | 0/2 | 0/2 | 1/6 | 0/1 |
| CSF1 | 0/1 | 0/2 | 0/1 | 0/3 | 1/2 | 0/2 | 0/2 | 0/1 | 0/3 | 1/1 | 0/4 | 0/2 | 0/5 | 0/1 | 1/1 | 0/2 | 0/2 | 2/6 | 1/1 |
| CCL2 | 0/1 | 0/2 | 0/1 | 0/3 | 0/2 | 0/2 | 0/2 | 0/1 | 0/3 | 1/1 | 0/4 | 0/2 | 1/5 | 0/1 | 1/1 | 0/2 | 1/2 | 2/6 | 1/1 |
| IL1B | 0/1 | 0/2 | 0/1 | 0/3 | 0/2 | 0/2 | 0/2 | 0/1 | 0/3 | 0/1 | 0/4 | 0/2 | 0/5 | 0/1 | 1/1 | 0/2 | 0/2 | 2/6 | 0/1 |
| IL6 | 0/1 | 1/2 | 1/1 | 0/3 | 0/2 | 0/2 | 0/2 | 1/1 | 2/3 | 1/1 | 1/4 | 1/2 | 2/5 | 0/1 | 1/1 | 0/2 | 0/2 | 1/6 | 0/1 |
| GCSF | 0/1 | 1/2 | 0/1 | 0/3 | 0/2 | 0/2 | 0/2 | 0/1 | 0/3 | 0/1 | 0/4 | 0/2 | 1/5 | 0/1 | 1/1 | 0/2 | 0/2 | 1/6 | 0/1 |
| CXCL10 | 0/1 | 1/2 | 1/1 | 0/3 | 0/2 | 0/2 | 0/2 | 0/1 | 0/3 | 1/1 | 1/4 | 1/2 | 1/5 | 0/1 | 0/1 | 0/2 | 0/2 | 1/6 | 0/1 |
| VEGFA | 0/1 | 0/2 | 0/1 | 0/3 | 0/2 | 0/2 | 0/2 | 1/1 | 0/3 | 0/1 | 0/4 | 0/2 | 0/5 | 0/1 | 0/1 | 0/2 | 0/2 | 2/6 | 1/1 |
| TNF | 0/1 | 0/2 | 0/1 | 0/3 | 0/2 | 0/2 | 0/2 | 0/1 | 0/3 | 0/1 | 2/4 | 0/2 | 1/5 | 0/1 | 0/1 | 0/2 | 0/2 | 2/6 | 1/1 |
| PDGFB | 0/1 | 0/2 | 0/1 | 0/3 | 0/2 | 0/2 | 0/2 | 0/1 | 0/3 | 0/1 | 0/4 | 0/2 | 0/5 | 0/1 | 1/1 | 0/2 | 0/2 | 1/6 | 0/1 |
| IL8 | 0/1 | 0/2 | 0/1 | 0/3 | 0/2 | 0/2 | 0/2 | 0/1 | 0/3 | 1/1 | 0/4 | 0/2 | 0/5 | 0/1 | 0/1 | 0/2 | 0/2 | 2/6 | 1/1 |
| PDGFA | 0/1 | 0/2 | 0/1 | 0/3 | 0/2 | 0/2 | 0/2 | 0/1 | 0/3 | 0/1 | 0/4 | 0/2 | 0/5 | 0/1 | 0/1 | 0/2 | 0/2 | 0/6 | 0/1 |
| IL12Bp40 | 0/1 | 1/2 | 0/1 | 0/3 | 0/2 | 0/2 | 0/2 | 0/1 | 0/3 | 0/1 | 0/4 | 0/2 | 0/5 | 0/1 | 1/1 | 0/2 | 0/2 | 0/6 | 0/1 |
| IL15 | 0/1 | 0/2 | 0/1 | 0/3 | 0/2 | 0/2 | 0/2 | 1/1 | 0/3 | 0/1 | 2/4 | 0/2 | 1/5 | 0/1 | 1/1 | 0/2 | 0/2 | 1/6 | 1/1 |
| CXCL1 | 0/1 | 0/2 | 0/1 | 0/3 | 0/2 | 0/2 | 0/2 | 0/1 | 0/3 | 0/1 | 0/4 | 0/2 | 1/5 | 0/1 | 0/1 | 0/2 | 0/2 | 2/6 | 1/1 |
| CCL4 | 0/1 | 0/2 | 0/1 | 1/3 | 0/2 | 0/2 | 0/2 | 0/1 | 0/3 | 0/1 | 0/4 | 0/2 | 1/5 | 0/1 | 0/1 | 0/2 | 0/2 | 0/6 | 0/1 |
| IFNA2 | 0/1 | 0/2 | 0/1 | 0/3 | 0/2 | 0/2 | 0/2 | 0/1 | 0/3 | 0/1 | 0/4 | 0/2 | 1/5 | 0/1 | 0/1 | 0/2 | 0/2 | 0/6 | 0/1 |
| IL13 | 0/1 | 0/2 | 0/1 | 0/3 | 0/2 | 0/2 | 1/2 | 0/1 | 0/3 | 0/1 | 0/4 | 0/2 | 1/5 | 0/1 | 0/1 | 0/2 | 0/2 | 2/6 | 0/1 |
| FLT3L | 0/1 | 0/2 | 1/1 | 0/3 | 1/2 | 0/2 | 0/2 | 0/1 | 0/3 | 0/1 | 0/4 | 1/2 | 0/5 | 0/1 | 1/1 | 0/2 | 1/2 | 0/6 | 0/1 |
| CD40LG | 0/1 | 0/2 | 0/1 | 0/3 | 0/2 | 0/2 | 1/2 | 0/1 | 0/3 | 0/1 | 0/4 | 0/2 | 0/5 | 0/1 | 1/1 | 0/2 | 0/2 | 0/6 | 0/1 |
| CCL22 | 0/1 | 0/2 | 1/1 | 0/3 | 0/2 | 0/2 | 0/2 | 1/1 | 0/3 | 0/1 | 0/4 | 0/2 | 1/5 | 0/1 | 1/1 | 0/2 | 0/2 | 2/6 | 1/1 |

Supplementary Figure S3. Detailed results of the association with different diseases of the immunologic markers studied. For each immunomarker, the number of disease sensitivity tests passed (Mann-Whitney p-value < 0.05) is given both overall and for each ICD-11 code.
